# Robust expression of LINE-1 retrotransposon encoded proteins in oral squamous cell carcinoma

**DOI:** 10.1101/2020.11.24.395285

**Authors:** Koel Mukherjee, Debpali Sur, Abhijeet Singh, Sandhya Rai, Neeladrisingha Das, Srinu Narindi, Rakshanya Sekar, Vandana Kumar Dhingra, Bhinyaram Jat, K V Vinu Balraam, Satya Prakash Agarwal, Prabhat Kumar Mandal

**Affiliations:** Department of Biotechnology, IIT Roorkee, Roorkee, Uttarakhand, India; Department of Head-Neck Surgery and Oncology, AIIMS Rishikesh, Rishikesh Uttarakhand, India; Armed Forces Medical College, Pune, India; School of Biosciences and Technology, Vellore Institute of Technology, Vellore, Tamil Nadu, India

**Keywords:** L1 retrotranspson, Cancer and L1 retrotransposon, L1ORF2p antibody, L1ORF1p antibody, OSCC and L1 retrotransposon

## Abstract

Retrotransposons are sequences which transpose within genomes using RNA as an intermediate. Long INterpersed Element-1 (LINE1 or L1) is the only active retrotransposon occupying around 17% of the human genome with an estimated 500,000 copies. An active L1 encodes two proteins (L1ORF1p and L1ORF2p); both of which are critical in the process of retrotransposition. In-order to propagate to the nextgeneration, L1s remain active in germ tissues and at an early stage of development. Surprisingly, by some unknown mechanism, L1 also shows activity in certain parts of the normal brain and many cancers. L1 activity is generally determined by assaying L1ORF1p because of its high expression and availability of the antibody. However, due to its lowerexpression and the unavailability of a robust antibody, detection of L1ORF2p has been limited. L1ORF2p is the crucial protein in the process of retrotransposition as it provides endonuclease and reverse transcriptase (RT) activity. Here, we report a novel human L1ORF2p antibody generated using an 80-amino-acid stretch from the RT domain, which is highly conserved among different species. The antibody detects significant L1ORF2p expression in murine germ tissues and human oral squamous cell carcinoma (OSCC) samples. This particular cancer is prevalent in India due to excessive use of tobacco. Here, using our in-house antibodies against L1 proteins, we show that more than fifty percent of samples are positive for L1 proteins. Overall, we reported a novel L1ORF2p antibody that detects L1 activity in germ tissues and OSCC

## 1. Introduction

Retrotransposons (jumping genes) are sequences which move from one place inthe genome to another using RNA as an intermediate, and areresponsible for individual cases of manygenetic disorder due to disruption of essential genes [1, 2, 3]. The human genome harbours a retrotransposon named Long INterspersed Element (LINE-1 or L1) which is highly active as evidenced by 500,000 copies occupying around 17% of the human genome [4, 5]. In addition to copying itself to new genomic locations, L1 activity is also responsible for the formation of over one million other retrotransposon insertions (e.g., Alu and SVA elements) and several thousand processed pseudogenes in the human genome [6, 7, 8, 9].

Although highly abundant, only a subset of L1s (~80-100 copies) are actively retrotransposing in any given human [10]. A retrotransposition-competent L1(RC-L1) is 6 kb in length with the following features: a 5’-UTR (~900 bp) with an internal promoter, two non-overlapping open reading frames (ORFs designated L1ORF1p and L1ORF2p) separated by a 63 bp spacer sequence, and a ~200 bp 3’-UTR ends with a poly (A) sequences with variable length (10-400 bp) [11, 12, 13]. The element is surrounded by target site duplications (TSDs) that vary both in size and sequence[2]. Human L1 ORF1 encodes a 40 kDa protein (338 amino acids in length) termed ORF1p comprised of three distinct domains; Coiled Coil (CC) (amino acids 52-153), RNA Recognition Motif (RRM) (amino acids 157-252), and Carboxy Terminal Domain (CTD) (amino acids 264-323) [14,15]. *In-vitro* studies have demonstrated that human ORF1p is a non-specific single-stranded nucleic acid-binding protein with nucleic acid chaperone activity [14, 15, 16]. Human ORF2 encodes a 150 kDa protein with reverse transcriptase (RT) [17] and endonuclease (EN) [18] activities.Functional studies have revealed that both ORF1p and ORF2p are critical for retrotransposition of their encoding RNA [13].

Due to their potential to function as insertional mutagens, L1s are generally silenced in somatic cells through epigenetic and post-transcriptional mechanisms. These include CpG methylation ofthe L1 promoter, small RNA induced silencing, cellular host factor-mediated retrotransposition inhibition and others [19, 20]. However, recent transgenic animal models and deep-sequencing studies revealed that a subset of L1s escaperepression and show high activity in germ cells, early stages of development, certain parts of the brain and in cancers [21–27].Thewhole-genome (WGS) and targeted sequencing of tumour samples showed high rates of L1 retrotransposition in many cancers,particularly those of epithelial cell origin [23]. Notably, in some cancers (e.g., colorectal cancer), the somatic L1 insertion frequency is striking with more than 100 retrotranspositionevents detected in one tumor [23], while in other tumors, retrotransposition is not found. Immunohistochemical analysis using human L1 ORF1p antibody showed that nearly half of human cancers are immune-reactive with LINE-1 ORF1p [25]. Recently, in a pilot study with a small number of samples, we have demonstrated significant expression of L1ORF1p in post-operative oral cancer, a cancer which is extremely common in India due to excessive use of tobacco [27]. The L1ORF2p is the central protein required in the process of retrotransposition and is very difficult to detect due to its low expression and lack of a specific antibody [13, 28–31]. Although our understanding of L1 activity in cancers has increased dramatically over the past five years, very few studies have been conducted to determinethe expression of L1 ORF2p in cancers due to lack of a proper ORF2p antibody [32–35].A few groups have reported an L1ORF2p antibody[32–35];however, questions have been raised regarding the specificity and sensitivity of those antibodies.

Here we reportthe successful development ofa human L1ORF2p antibody using an 80aminoacid stretch from the reverse transcriptase domain of L1ORF2p. Alignment of this human stretch with mouse and rat L1ORF2p sequences showed that the stretch is highly conserved among all three species [36]. The developed L1ORF2p antibody detects significant ORF2p expression in murine testis and ovaryand in post-operative oral cancer samples. Until now, no study has reported L1ORF2p expression in oral cancer. We have screened 39 post-operative oral cancer samples and found that nearly fifty percent of samples expressed L1ORF2p. Overall, we reported a novel ORF2p antibody which detects L1 activity in germ tissues and post-operativeoral cancer samples.

## 2. Materials and Methods

### 2.1. Cloning of human hL1RT_EH_ fragment

The human L1 ORF2pRT domain fragment (234 bp*EcoRI -HindIII* fragment, Nucleotide position in L1ORF2 1435-1674, amino acid position in L1ORF2 479-558, L1RP accession number AF148856) [37] was cloned in pET28a to make a pEThL1RT_EH_ clone. The cloning strategy is provided in Figure 1 and Supplementary Figure 1.

**Figure 1:**
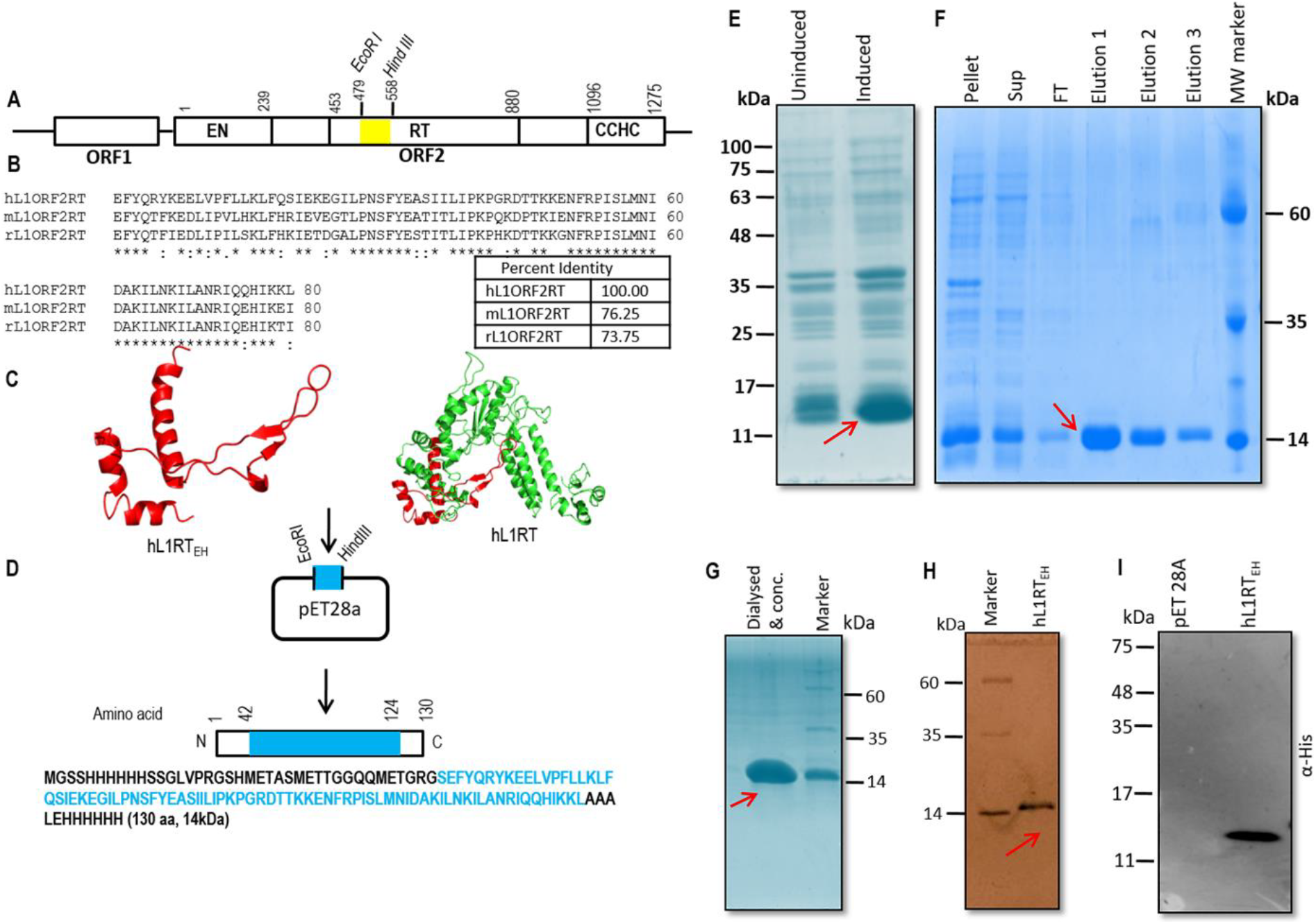
Generation of the antigen for the production of Human L1 ORF2 specific antibody. (A) Schematic diagram of full length active human L1 retrotransposon with two encoded proteins (L1ORF1p and L1ORF2p). L1ORF2p (1275 amino acids in length with predicted MW 150kDa) has three partially characterized domains: endonuclease (EN) (AA:1-239), reverse transcriptase (RT) (AA: 453-883) and a cysteine-histidine-rich domain (CCHC) (AA: 1096-1275). The *EcoRI-HindIII* restriction fragment (AA: 479-558 fragment name hL1RT_EH_) from the RT region was used as an antigen to make antibody against hL1ORF2p (shaded by yellow). (B) Alignment of the selected eighty amino acids stretches of the RT domain (hL1RT_EH_)among human, mouse and rat L1. The selected RT stretch showed 76.25% and 73.75% identity at the protein level with the same stretch present in mice and rat L1ORF2, respectively. (C) Predicted structure of hL1RT_EH_ fragment and complete human RT domain (generated using PyMOL) [58].(D) Sub-cloning scheme of human hL1RT_EH_fragment in pET-28a bacterial expression vector. Bacterial expressed hL1RT_EH_ fragment encodes 130 amino acid polypeptide (from N terminal to C terminal AA: 1-42 vector, 43-124 RT fragment, and 125-130 vector) with predicted MW around 14 kDa. (E) Whole-cell lysate SDS-PAGE of *E.coli* expressed pet-hL1RT_EH_. Induced protein with a molecular weight around 14 kDa is shown by the arrow. (F) Purification of hL1RT_EH_ from inclusion bodies by dissolving the pellet fraction in 8M urea buffer (details in materials and methods). The purified protein in elution 1, 2 and 3 is shown by an arrow. (G) Dialysed and concentrated hL1RT_EH_fragment protein (antigen) injected to mice for antibody generation. (H) Silver staining of purified hL1RT_EH_ (antigen) show the purity of antigen used to generate the antibody. (I) Western blotting of hL1RT_EH_ using an anti-His antibody. MW: Molecular Weight, FT: Flow-through.

### 2.2. Expression and Purification of Human hL1RT_EH_ domain fragment protein

The pEThL1RT_EH_clone was transformed to *E.coli*(strain BL21) expression cells, plated to Kanamycin containing agar plates and incubated at 37°C overnight. A single colony was incubated overnight in 10 ml LB media with an appropriate antibiotic (Kanamycin 25 μg/ml) to obtain primary culture. One percent of the primary culture was added to 100 ml LB media and the culture was grown at 37°C till OD_600_ 0.4 before the addition of 0.4mM isopropyl β-D-thiogalactopyranoside (IPTG) for the induction of hL1RT_EH_at 37°C for another 3 hours. After the induction, cells were harvested, resuspended in lysis buffer [50mM Tris-cl (pH-8.0), 150mM NaCl, 10mM imidazole, 1mM PMSF]. The cells were then lysed by three cycles of freeze-thawing followed by sonication on ice.The lysate was centrifuged at 12000 x g for 30 minutes at 4°C. As the hL1RT_EH_ formed inclusion bodies, the supernatant was discarded, and the pellet was dissolved in buffer A (50 mM Tris-Cl pH 8.0, 150 mM NaCl, 10 mM imidazole, 8M Urea) followed by centrifugation at 12000 x g for 20 minutes at 4°C. The supernatant was collected to a separate tube, and the pellet was discarded. Next, the supernatant was incubated with pre-equilibrated nickel agarose resin (Qiagen) (100 ul packed resin per 50 ml culture) for one hour at 4°C with continuous shaking. The resin was washed with 5 ml wash buffer (50mM imidazole).The protein was finally eluted in elution buffer (50 mM Tris-Cl pH 8.0, 150 mMKCl, 300mM imidazole, 6M Urea). The urea was by removed serial dialysis to avoid precipitation of protein, i.e. 8M Urea→6M Urea→ 4M Urea→ 2M Urea→ 1M Urea→ 0.8M Urea→ 0.6M→0.4M→0.2M→0M Urea.After dialysis, the proteinwas concentrated using a centrifugal filter unit (MWCO 3000 Da) and stored at −20°C after a flash freeze. Purified protein fragment with a molecular mass of approximately 14 kDa (vector sequence plus eighty amino acid RT fragment, details in Figure 1 and supplementary text) was used to immunize mice (Swiss Albino).

### 2.3 Generation of polyclonal antibody against hL1RT_EH_ in mice

The purified hL1RT_EH_ protein was used as an antigen for raising its antibody in swiss albino mouse. The 80 days immunization protocol followed is discussed below in details:

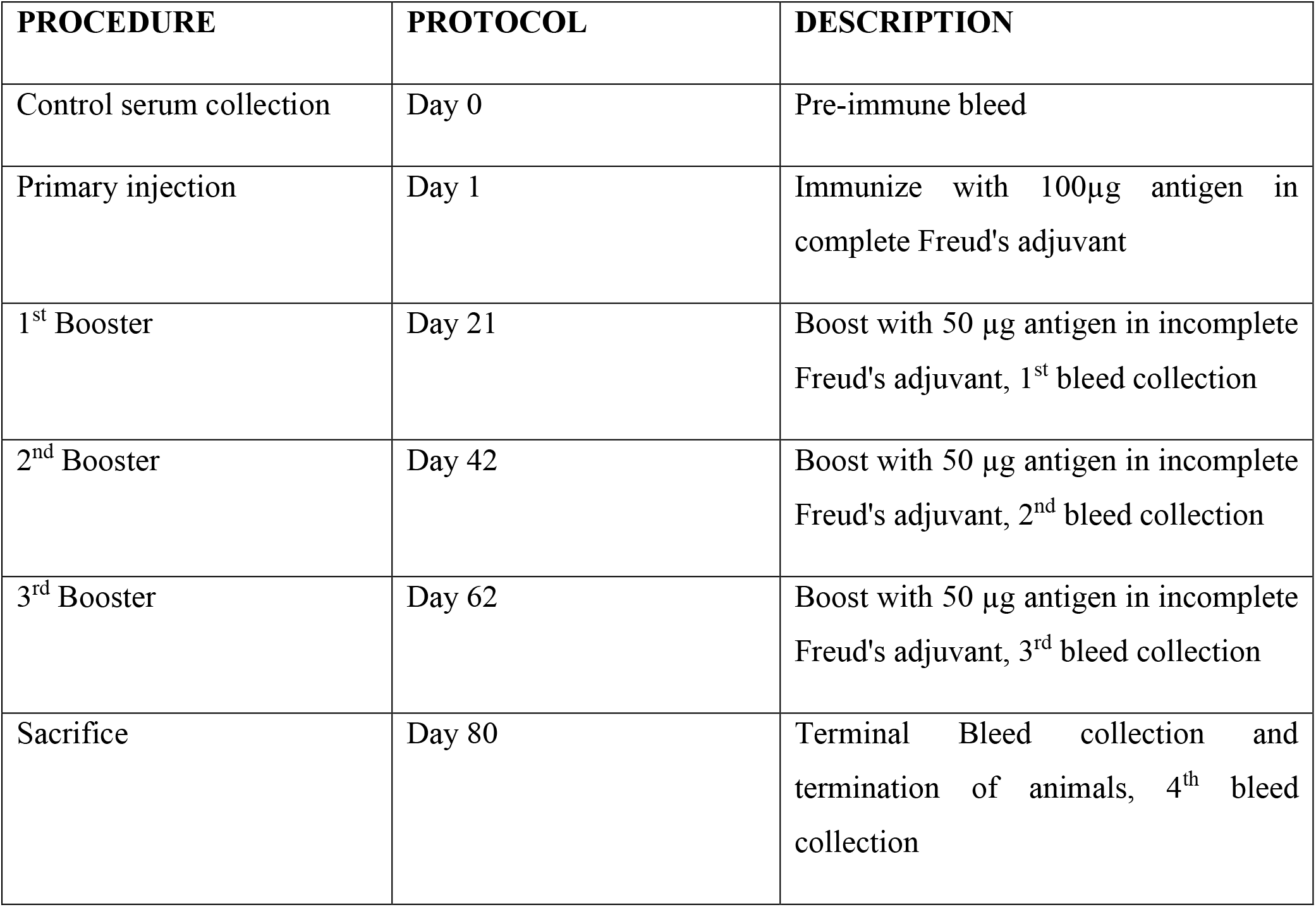

| PROCEDURE | PROTOCOL | DESCRIPTION |
| --- | --- | --- |
| Control serum collection | Day 0 | Pre-immune bleed |
| Primary injection | Day 1 | Immunize with 100µg antigen in complete Freud's adjuvant |
| 1<sup>st</sup> Booster | Day 21 | Boost with 50 µg antigen in incomplete Freud's adjuvant, 1<sup>st</sup> bleed collection |
| 2<sup>nd</sup> Booster | Day 42 | Boost with 50 µg antigen in incomplete Freud's adjuvant, 2<sup>nd</sup> bleed collection |
| 3<sup>rd</sup> Booster | Day 62 | Boost with 50 µg antigen in incomplete Freud's adjuvant, 3<sup>rd</sup> bleed collection |
| Sacrifice | Day 80 | Terminal Bleed collection and termination of animals, 4<sup>th</sup> bleed collection |

Serum was isolated from blood collected in each step by incubating the blood for 1 hour, in 37°C water bath followed by incubating a 4°C over-night. Next day the sample was centrifuged (12,000xg, 10 minutes, 4°C), and the serum was collected.

### 2.4. Purification of IgG fraction of anti-hL1RT_EH_ from whole immunized serum

Around 500μl of total serum obtained from hL1RT_EH_ fragment immunized mice was mixed in 1:1 ratio with binding buffer (0.02M sodium phosphate, pH 7).Around100μl packed protein G agarose slurry (stored in 20% ethanol) was taken and equilibrated with binding buffer. Next, serum and binding buffer mix (ratio1:1, Volume 1000 μl) was added to the beads and incubated for 1 hour at 4°C (in rocker) for effective binding of IgG with protein G agarose.After incubation centrifuged the tube at 2000 xg for one min at 4°C, and incubated the tube on ice for one minute to settle the beads. The flow-through was carefully removed and stored for further analysis. Next, the beads washed with 5 x 1 ml wash buffer (0.02M sodium phosphate, pH 7). Finally, IgG was eluted in 100 μl elution buffer (0.1M Glycine, pH 2.5) by centrifugation at 2000 x g for 2 min at 4°C. Immediately the eluate was neutralized with 23 μl of neutralization buffer (1M Tris-Cl, pH-9.0).The concentration of the pure antibody was carried out using Bradford assay, and the quality of the purified antibody was checked in 10% SDS-PAGE gel. The affinity-purified antibody was checked and confirmed by performing immunoblotting.

### 2.5. Animals

The Swiss albino mice and Sprague Dawley rats were procured from the animal facility of National Institute of Pharmaceutical Education and Research (NIPER), Chandigarh, India were housed in the animal facility at Indian Institute of Technology Roorkee, India. All the experimens were carried out as per indicated guidelines of the Institute Animal Ethics Committee (IAEC) underProtocol no: BT/IAEC/2018/12.

### 2.6. Cell culture

HEK293T, (human embryonic kidney) and HeLa (cervical carcinoma) cells obtained from National Centre for Cell Science (NCCS), Pune, India were maintained in a CO_2_ incubator at 37°C and 5% CO_2_ concentration in high glucose Dulbecco’s modified Eagle medium (DMEM) with L– glutamine (Gibco) supplemented with 10% fetal bovine Calf serum and 100 U/ml penicillin-streptomycin (Himedia).

### 2.7. Plasmids and cell Transfection

Around 2 ×10^5^ HEK293T (human embryonic kidney) were seeded on a 35 mm tissue culture plate. After 8-10 hours incubation, 1μg of L1-EGFP construct was transfected into cells using Fugene 6 Transfection Reagent. One day after transfection the old media was replaced with new media; 96 hours after post-transfection EGFPpositive cells were checked under the microscope to proceed for immune-blotting.

### 2.8. Cells and tissue lysate preparation and immunoblotting

Whole-cell lysates from cell lines were prepared using NP40 lysis buffer [20 mM Tris-Cl pH 7.8‚137 mM NaCl and 1% NP-40 supplemented with 1X protease inhibitor cocktail (Roche)].The tissue lysate was prepared from mouse, rat and OSCC tissues using RIPA buffer [50mM Tris-Cl pH 8, 150mM NaCl, 0.5% sodium deoxycholate, 0.1% SDS and 1% NP-40 supplemented with 1X protease inhibitor cocktail (Roche)]. The lysate was centrifuged at 10,000 g for 10 minutes at 4°C, and the supernatant was transferred to a new 1.5 ml tube and stored at −70°C until further use. Around 40 μg of whole-cell lysate was resolved in 10% SDS-PAGE gel (Mini protein Tetra cell Bio-Rad) and blotted to nitrocellulose membrane (Bio-Rad) by applying 100V for 75 minutes using Bio-Rad mini trans blot electrophoretic transfer cell (transfer buffer composition:25mM Tris-Cl pH 7.6, 192mM Glycine, 0.1% SDS, 20% Methanol). The membrane was probed with Primary: polyclonal mouse anti-hL1RT_EH_ (1:5000),anti-hL1-ORF1 (RRM) (1:33000) [Sur et al., 2017], anti-GAPDH (1:6000) (Santacruz Biotechnology). The next day, the membrane was washed for 1hour using 1x TBS-T (5×2, 10×2, 15×2min), incubated with conjugated secondary α-rabbit HRP and secondary α-mouse HRP (Jacksons Immuno Research Laboratories, USA). Western blots were developed using ECL western blotting detection reagent (Biorad) as per the manufacturer’s instructions. The bands were detected by exposing the blot on X-ray film (Biorad).

### 2.9. Sample collection of cancer tissue specimens

The Indian Institute of Technology Roorkee (IITR) and All India Institute of Medical Sciences (AIIMS) Rishikesh have signed Memorandum of Understanding (MOU) for conducting joint research on patient samples as per Institute ethical guidelines. Post-operative cancer tissues were collected from the surgical oncology department AIIMS Rishikesh following proper written consent from the patient and their immediate family member as per institute guidelines. The patient and/or the family members understood and agreed that a small portion (2-5 gram) of operated cancer tissue will be used for research purpose to understand the biology of oral cancer and its treatment. Clinical details of patients used in this study are present in (Table 1). Following initial collection, samples were stored in RNA later solution (Qiagen) at −20°C and used for lysate preparation. The tissue samples stored in 10% NBF solution were subsequently used for the generation of formalin-fixed paraffin embedded blocks (FFPE). All the experiments were conducted in accordance with ethical principles embodied in the declaration of tissue request and material transfer agreement between AIIMS Rishikesh and IIT Roorkee. The approval from the institutional ethics committee of All India Institute of Medical Sciences, Rishikesh (Reg No: ECR/736/Inst/UK/2015/RR-18) has been obtained specifically for the work with human samples (Letter No: AIIMS/IEC/20/395). All the investigations in this study strictly followed the rules set by the Declaration of Helsinki.

**Table 1:** The details of patients used in this study.

| Sl.No. | Age | Sex | Site of Tumor | Grade | P TNM | L1 ORF1p | p53 | L1 ORF2p |
| --- | --- | --- | --- | --- | --- | --- | --- | --- |
| C1 | 39 | M | Left upper alveolus | Moderately differentiated squamous cell carcinoma | T4A N1 | High | Negative | High |
| C2 | 56 | F | Right buccal mucosa | Well differentiated squamous cell carcinoma | T2 N0 | High | Positive | High |
| C3 | 27 | M | Left buccal mucosa | Well differentiated squamous cell carcinoma | T4A N2B | Negative | Negative | Negative |
| C4 | 38 | M | Tongue | Moderately differentiated squamous cell carcinoma | T2 N2C | Negative | Positive | Negative |
| C5 | 33 | M | Right buccal mucosa | Moderately differentiated squamous cell carcinoma | T4A N2B | Negative | Positive | Negative |
| C6 | 60 | M | Left buccal mucosa | Well differentiated squamous cell carcinoma | T2 N0 | High | Negative | High |
| C7 | 45 | M | Left buccal mucosa | Well differentiated squamous cell carcinoma | T3 N2B | Negative | Negative | Negative |
| C8 | 75 | F | Left buccal mucosa | Well differentiated squamous cell carcinoma | T4A N0 | Moderate | Negative | Moderate |
| C9 | 50 |  | Right buccal mucosa | Moderately differentiated squamous cell carcinoma | T4A N2B | Moderate | Positive | Moderate |
| C10 | 33 | M | Left buccal mucosa | Moderately differentiated squamous cell carcinoma | T4A N0 | Negative | Negative | Negative |
| C11 | 28 | F | Tongue | Poorly differentiated squamous cell carcinoma | T2N2C | Moderate | Negative | Moderate |
| C12 | 40 | M | Left buccal mucosa | Well differentiated squamous cell carcinoma | T3 N0 | Negative | Negative | Negative |
| C13 | 43 | M | Left buccal mucosa | Moderately differentiated squamous cell carcinoma | T4A N2B | Negative | Negative | Negative |
| C14 | 38 | M | Right upper alveolus | Well differentiated squamous cell carcinoma | T4A N0 | Negative | Negative | Negative |
| C15 | 37 | M | Left tongue | Moderately differentiated squamous cell carcinoma | T4aN2Mx | Moderate | Positive | Moderate |
| C16 | 42 | M | Left buccal mucosa | Moderately differentiated squamous cell carcinoma | T4A N1 | Negative | Negative | Negative |
| C17 | 28 | M | Lower lip | Moderately differentiated squamous cell carcinoma | T2N2C | Moderate | Negative | Moderate |
| C18 | 60 | M | Right tongue | Moderately differentiated squamous cell carcinoma | T2 N2B | High | Negative | High |
| C19 | 46 | M | Left buccal mucosa | Moderately differentiated squamous cell carcinoma | T4A N0 | High | Positive | High |
| C20 | 38 | M | Right tongue | Well Differentiated Squamous Cell Carcinoma | T4A N1 | High | Positive | High |
| C21 | 50 | M | Right buccal mucosa | Moderately differentiated squamous cell carcinoma | T2 N0 | Negative | Positive | Negative |
| C22 | 34 | M | Right buccal mucosa | Moderately differentiated squamous cell carcinoma | T3 N0 | High | Negative | High |
| C23 | 40 | M | Right buccal mucosa | Moderately Differentiated Squamous Cell Carcinoma | T4A N0 | High | Positive | High |
| C24 | 40 | M | Left buccal mucosa | Well differentiated squamous cell carcinoma | T4A N2B | High | Positive | High |
| C25 | 47 | M | Right buccal mucosa | Moderately Differentiated Squamous Cell Carcinoma | T4A N0 | Negative | Negative | Negative |
| C26 | 37 | M | Right buccal mucosa | Moderately Differentiated Squamous Cell Carcinoma | T3N2B | Negative | Positive | Negative |
| C27 | 59 | F | Left buccal mucosa | Moderately differentiated squamous cell carcinoma | T4A N0 | Negative | Negative | Negative |
| C28 | 40 | M | Left buccal mucosa | Moderately differentiated squamous cell carcinoma | T4A N2B | Moderate | Negative | Moderate |
| C29 | 51 | M | Right buccal mucosa | Moderately differentiated squamous cell carcinoma | T2N0 | Negative | Negative | Negative |
| C30 | 50 | M | Left buccal mucosa | Poorly differentiated squamous cell carcinoma | T4A N2B | Negative | Negative | Negative |
| C31 | 50 | M | Lower lip | Moderately differentiated squamous cell carcinoma | T4A N2C | Negative | Negative | Negative |
| C32 | 44 | M | Left buccal mucosa | Moderately differentiated squamous cell carcinoma | T4AN0 | High | Positive | High |
| C33 | 52 | M | Tongue | Well differentiated squamous cell carcinoma | T4AN2B | High | Positive | High |
| C34 | 40 | M | Left buccal mucosa | Moderately differentiated squamous cell carcinoma | T2N0 | Negative | Negative | Negative |
| C35 | 34 | M | Right buccal mucosa | Moderately differentiated squamous cell carcinoma | T4AN0 | High | Positive | High |
| C36 | 43 | M | Right buccal mucosa | Moderately differentiated squamous cell carcinoma | T1N0Mx | Negative | Negative | Negative |
| C37 | 44 | M | Tongue | Moderately differentiated squamous cell carcinoma | T3N2M<sub>x</sub> | High | Negative | High |
| C38 | 43 | M | Right buccal mucosa | Moderately differentiated squamous cell carcinoma | T4AN1 | Moderate | Negative | Moderate |
| C39 | 46 | M | Left buccal mucosa | Moderately differentiated squamous cell carcinoma | T3N2B | Negative | Positive | Negative |

### 2.10. Hematoxylin-eosin staining

Formalin-fixed paraffin-embedded (FFPE) tissue sections of 4 microns were deparaffinised using xylene followed by rehydration in descending grade of ethanol solutions. Then the slides were stained with Mayer’s hematoxylin (Himedia) for three minutes, washed in tap water and counterstained with eosin Y solution (Himedia) for one minute. The slides were further dehydrated with ascending grade of ethanol solutions, and finally, the tissue sections were examined under an upright light microscope (Leica Microsystems) after mounting with DPX mounting media (Himedia).

### 2.11. Immunohistochemistry

Formalin-fixed paraffin-embedded (FFPE) tissues weresectioned to4 micron thickness on coated glass slides. They were deparaffinized, and rehydrated in descending grades of ethanol solutions before proceeding for antigen retrieval. The antigen retrieval was performed in a common household vegetable steamer (pressure cooker) using Tris-EDTA antigen retrieval buffer (10 mM Tris base, 1 mM EDTA solution, 0.05% Tween 20, pH-9.0), following it slides were washed 2 × 5 minutes each in TBST buffer (1X TBS containing 0.025% Triton-X100) and then blocked in blocking solution (1% BSA in 1XTBST) for 1 hour at room temperature. For ORF1p staining, s?lides were incubated with polyclonal rabbit α-ORF1p (RRM) antibody (1:500 diluted in blocking solution), and for ORF2p we probed the slides with polyclonal mouse anti-hL1RT_EH_ (1:200 diluted in blocking reagent) and kept at 4°C overnight in a humid chamber. The next day, slides were washed with 1XTBST for 3x 10 min each with gentle agitation to remove unbound primary antibodies. Endogenous peroxidase activity was quenched by treating the slides with 0.3% hydrogen peroxide. Slides were then incubated with secondary antibody (1:500 dilution α-rabbit HRP and 1:500 dilution α-mouse HRP (Jacksons Immuno Research)) for an hour at room temperature. The slides were washed again 3 x 10 minutes with 1X TBST at room temperature with gentle agitation. All the signals were visualized by adding 3-3′-Diaaminobenzidinetetrahydrochloride (DAB substrate) solution (Roche) to the slides and counterstained with haematoxylin. (Himedia) Following counterstaining, the slides are dehydrated with ascending order of ethanol and mounted with DPX mounting media (Himedia). Anti-GAPDH (1:250) (Santacruz Biotechnology) was used as housekeeping control, PanCK (pre-diluted from PathnSitu biotechnology India) and Ki67((pre-diluted from PathnSitu biotechnology India)) were used as squamous cell carcinoma marker and cell proliferation marker respectively. Images were captured using an upright light microscope (Leica Microsystems) equipped with a camera. All the microscopic pictures were taken at 40X magnification.

## 3. Results

### 3.1. Generation of a novel polyclonal antibody against L1-ORF2p

Human L1 ORF2p protein is 1275 amino acid residues in length (L1RP, Accession number: AF148856.1) with a predicted molecular weight (MW) of~150 kDa. It has three partially characterized domains, which are from N-C terminal: endonuclease (EN) (AA:1-239), reverse transcriptase (RT) (AA: 453-883) and a cysteine/histidine-rich domain (CCHC) (AA: 1096-1275) [13] (Figure 1A). The L1ORF2p is the central protein required in the process of retrotransposition with demonstrated RT and EN activities[17,18].Due to its very low expression, the detection of ORF2p is extremely challenging[28–31]. Deciphering the expression level of human L1-ORF2p across various somatic, germline and cancerous tissues could lead to a better understanding of L1 biology.

In this study, we have generated a highly specific polyclonal antibody against human L1-ORF2. Through bioinformatics analysis, we identified an 80 amino acid stretch present in the RT domain [240bp*EcoRI-HindIII* fragment of L1ORF2, Nucleotide position in L1ORF2 1435-1674, amino acid position in L1ORF2 479-558, (L1RP accession number AF148856)], to be evolutionarily conserved across various species (Figure 1A and 1B). Multiple sequence alignment results showed that the selected RT stretch is 76.25% and 73.75% identical at the protein level with mouseand rat L1 RT, respectively (Figure 1B and supplementary text). The *in slico* structural study showed that the selected stretch is open and situated outside of the folded RT structure; thus, it can be a suitable epitope for antibody generation (Figure 1C).

As the selected region is flanked by natural restriction enzymes *EcoRI* and *HindIII* in L1RP[37], the JCC5 clone (L1RP cloned inpBS) was digested with *EcoRI* and *HindIII* to get the desired fragment (234 nucleotides) and then sub-cloned into abacterial expression vector pET-28A (Figure 1A, Supplementary figure 1A). The resultant clone (named hL1RT_EH_), if expressed in a bacterial cell, will produce a 130 amino acid polypeptide (from N terminal to C terminal AA: 1-42 vector, 43-124 RT fragment, and 125-130 vector) with MW around 14 kDa (Figure 1D). The N-terminal 6X His-tagged hL1RT_EH_was expressed in *E. coli* expression cells, and a distinct band of around 14 kDa was detected in the pellet fraction, suggesting that the induced protein formed inclusion bodies (Figure 1E and 1F). The protein was then purified from inclusion bodies by dissolving the pellet in urea, followed by Ni-agarose chromatography (Figure 1F). Analysis of the purified hL1RTR_EH_fragment bySDS-PAGE followed by Coomassie and silver staining revealed a distinct band at around 14 kDa with ~100% purity (Figure 1G and 1H). The purified hL1RT_EH_ was also confirmed by Western blotting using an anti-His antibody before injecting to mouse (swiss albino) for antibody production (Figure 1I).The mouse preimmune serum was checked by Western blotting, and no cross-reactivity was detected (Supplementary Figure1B).

### 3.2. Characterization of α-hL1-ORF2 by immunoblotting

The mice anti-serum raised against hL1RT fragment was checked by Western blotting using IPTG-induced total lysate obtained from bacterial expression cells containing the pET-hL1RT_EH_clone (Figure 2A). A single band at around 14 kDa was detected in that lane, and the control lanes lackedany signals [total lysate from induced empty vector pET28a and pET-hRRM clone; ORF1p RRM domain cloned in pET30b [26] (Figure 2A).This resultsuggests that the selected ORF2p RT fragment (amino acids 479-558 of L1ORF2p) is immunogenic in mice and doesnotcross-react with human L1ORF1p protein (Figure 2A). Recently, we have reported a novel hL1ORF1 antibody where we use the RRM domain of human L1ORF1p as an antigen[26, 27]. Inorder to check whether anti-hL1ORF1 cross-reacts with thehL1ORF2pRT_EH_ fragment, a Western blot was performed with bacterial over-expressed pET-hRT_EH_, and no band was detected, suggesting that anti-hL1ORF1p doesnotcross-react with hORF2RT_EH_ peptide (Figure 2B).

**Figure 2:**
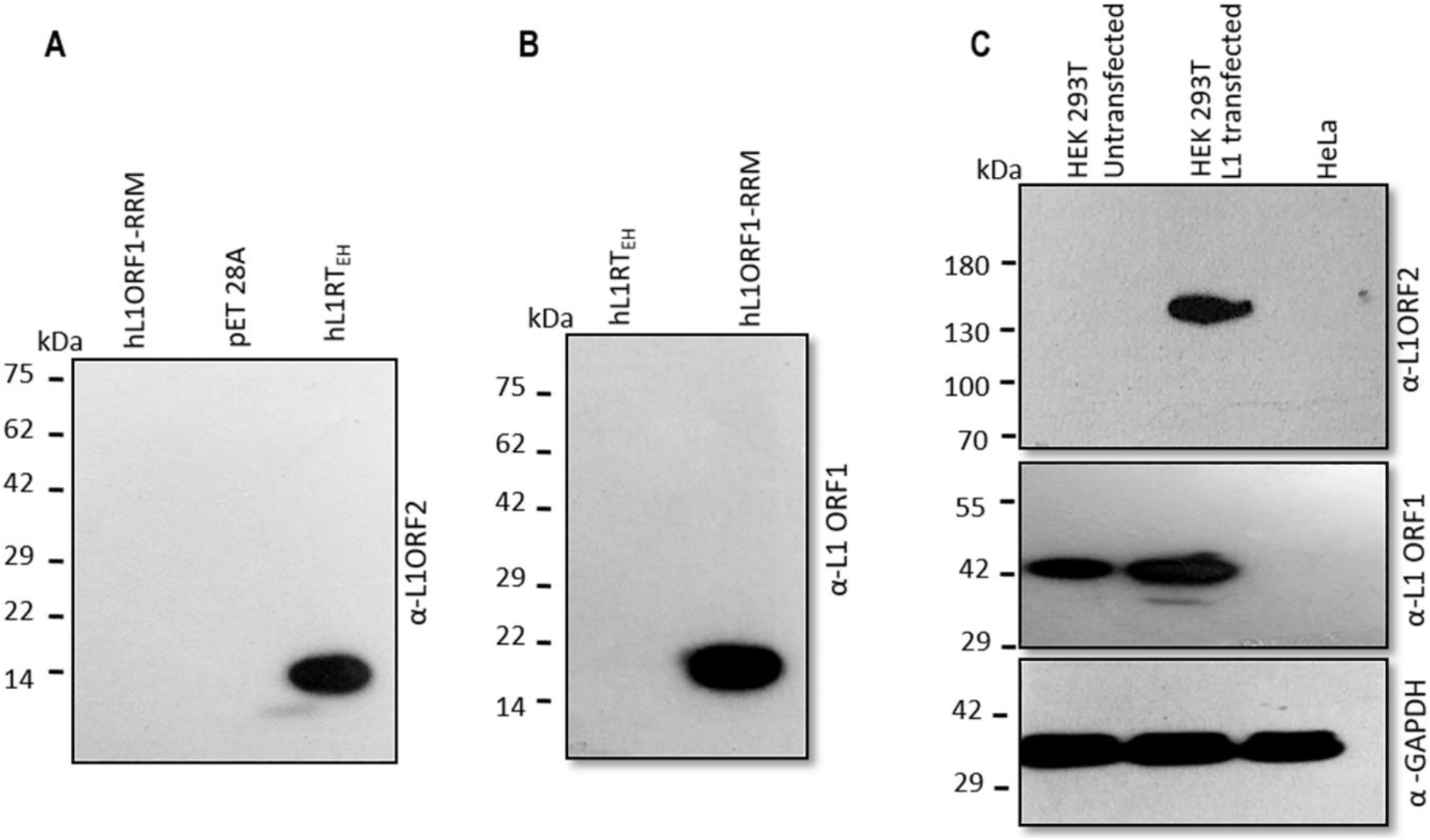
Characterization of human L1 ORF2p antibody (α-hL1ORF2) (A)Immunoblot analysis with anti-hL1ORF2RT_EH_ on induced bacterial lysate obtained from and pEThL1ORF1_RRM_, pET28A and pET-hL1RT_EH_. Human L1ORF1p RRM antigen used to make anti-hL1ORF1p didn’t show any cross-reaction with anti-hL1ORF2p [26,27] (B) Immunoblot analysis with anti-hL1ORF1_RRM_ on total bacterial lysate obtained from pET-hL1RT_EH_, and pEThL1ORF1_RRM_. No cross-reaction of anti-hL1ORF1with hL1RT_EH_ fragment used as antigen to make ORF2p antibody (Sur et al., 2017). (C) Detection of L1ORF2p (exogenous), L1ORF1p (endogenous and exogenous) and GAPDH (endogenous) [26,27] by immunoblotting in HEK 293T cells after transfecting full-length L1-EGFP construct (Ostertag 99RPEGFP).

Next,to test the anti-hRT_EH_antibody for its ability to detect native ORF2p in situ, we transfected episomal retrotransposon reporter plasmid 99RPEGFP (full-length disease-causing L1 with retrotransposon indicator cassette cloned in a pCEP4 vector) [38] into HEK293T cells. We performed Western blotting using total lysate 96 hours post-transfection. The data showed a distinct band at around 150 kDa, corresponsing to the proposed MW of ORF2p in the transfected lane, whereas no signal was detected in lysate obtained from untransfected cells (Figure 2C and Supplementary Figure 2A). The same samples were checked for the presence of hORF1p, and the Western blot showed the presence of ORF1p in both transfected and untransfected cells (endogenous ORF1p) (Figure 2C). Previous studies showed a significant amount of endogenous ORF1p expression in HEK293T cells[27]. GAPDH was used as a loading control (Figure 2C). From the above result, it is evident that the generated antibodyishighly specific and detects L1ORF2p as a discrete single band.

Next, to check the sensitivity of anti-ORF2p antibody, bacterial over-expressed pET-hORF2RT_EH_and 99RPEGFP [38] transfected HEK293T total lysate were separated in SDS-PAGE gel in increasing concentration, and Western blotting was performed. The data showed that the anti-ORF2p could detect as little as 10 ng and 20 μg in over-expressed bacterial and L1 transfected HEK293T total lysate, respectively (Supplementary Figure 2B).

### 3.3. Detection of endogenous L1ORF2p and L1ORF1p in different somatic and germline tissues

It is well known that L1 retrotransposon activity is generally attenuated in somatic cells/tissues, while numerous reports indicate high L1 activity in germline cells and tissues [22,29,39].So, we wished to validate the above fact by evaluating the endogenous expression of L1ORF2 and L1ORF1 proteins in somatic and germline tissues with the generated antibodies. The amino acid stretch from the RT domain used to generate antibody showed significant similarity between human, mice and rat (Figure 1B). It indicates that the antibody generated using human RT fragment can detect L1ORF2p in mouseand rat. Hence, we wanted to explore the endogenous expression of L1ORF2p in mouse and rat tissues. Tissue lysates from kidney, liver, ovary and testis were prepared, and Western blotting was performed with anti-hL1RT_EH_ antibody for L1ORF2p detection (Figure 3A and 3B, upper panel). As expected, the somatic tissues of mice and rat (kidney and liver) did not show any expression of L1ORF2p. In contrast, the germ tissues (ovary and testis) showed robust expression of L1ORF2 protein at around 150 kDa (Figure 3A and 3B, upper panel). All the tissues were also assayedfor L1ORF1p expression; the mice and rat germ tissues (testis and ovary) showed significant expression of ORF1p (Figure 3A and 3B, middle panel), whereas no expression was detected in somatic tissues (kidney and liver) (Figure 3A and 3B, middle panel). GAPDH was used as a loading control for bothmouse and rat tissues (Figure 3A and 3B, lower panel).

**Figure 3:**
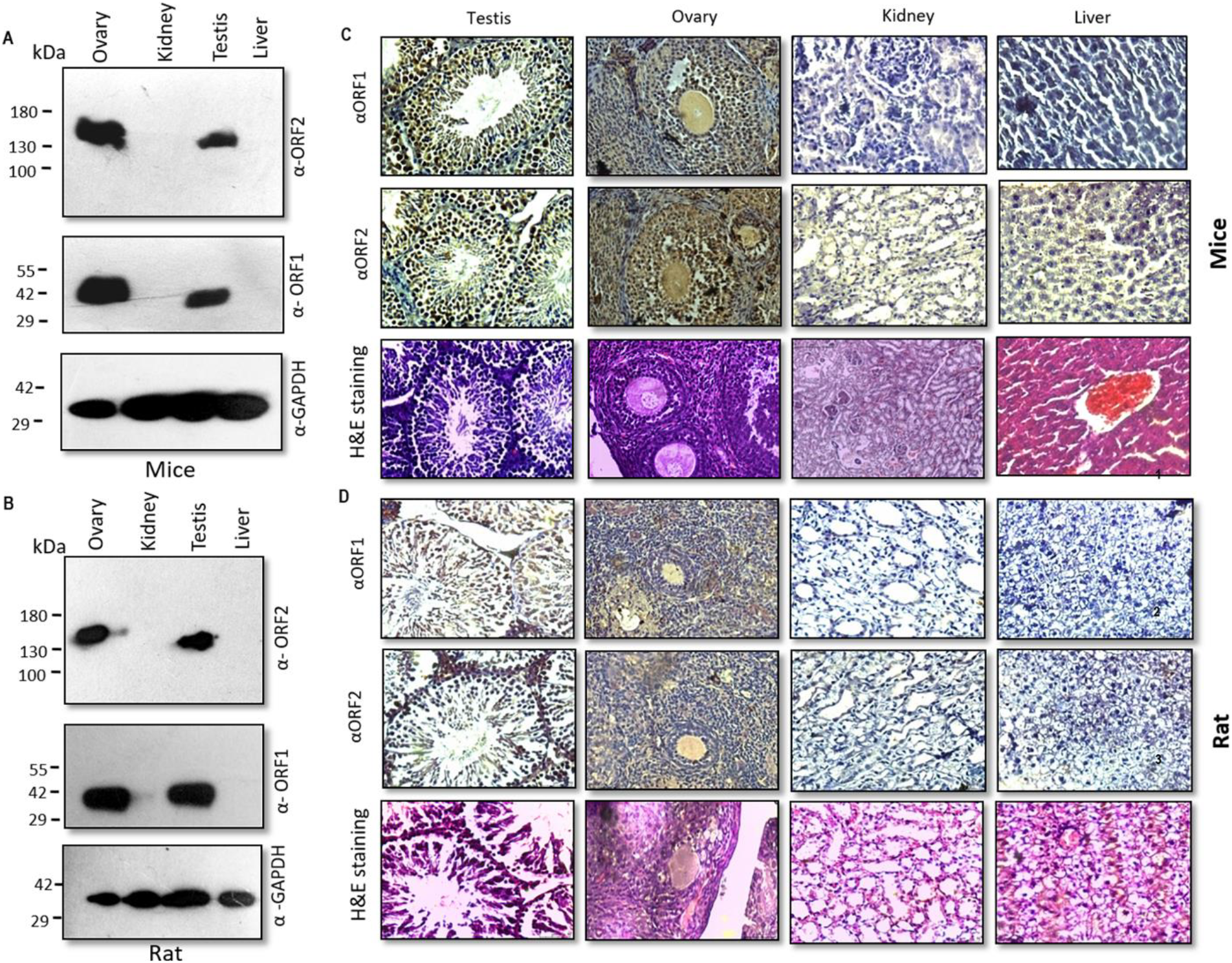
Detection of L1ORF1p and L1ORF2p in mouse and rat tissues. (A) Immunoblot analysisof somatic and germline tissues of the mouse (testis, ovary, liver and kidney) with anti-L1ORF2 (panel 1), anti-L1ORF1 (panel 2), and anti-GAPDH (panel 3) as the loading control. (B) Immunoblot analysis of somatic and germline tissues of rat (testis, ovary, liver and kidney) with anti-L1ORF2 (panel 1), anti-L1ORF1 (panel 2), and GAPDH (panel 3) as loading control. (C) Immunohistochemicalanalysis of somatic and germline tissues of the mouse (testis, ovary, liver and kidney) with anti-L1ORF1 (panel 1) and anti-L1ORF2 (panel 2). Tissue sections stained with hematoxylin-eosin are shown in panel 3. (D) Immunohistochemical analysis of somatic and germline tissues of rat (testis, ovary, liver and kidney) with anti-L1ORF1 (panel 1) and anti-L1ORF2 (panel 2). Hematoxylin-eosin stained samples are shown in panel 3.

Next, we performed immunohistochemistry (IHC) to complement our Western blotting result as well as to detect which group of cells express L1ORF2p protein in mice and rat (Figure 3C and 3D). Kidney, liver, testis and ovary tissues were selected both from mouse and rat to see the expression of ORF2p via IHC.The morphology of all the tissue sections was first checked by eosin hematoxyline staining (Figure 3C and Figure 3D, upper panel). The IHC analysis showed that both testis and ovary are expressing significant amounts of LIORF2p (Figure 3C and Figure 3D, lower panel). All the tissues were also checked for ORF1p expression; the mice and rat germ tissues (testis and ovary) showed significant expression of ORF1p (Figure 3C and Figure 3D, middle panel), whereas no or significantly less expression was detected in somatic tissues (kidney and liver) (Figure 3A and 3B).

### 3.4. Detection of L1ORF2p in OSCC

We next examined the L1ORF2p expression in post-operative OSCC samples. Recently,in a pilot study with a limited number of samples (n=15),we showed significant expression of L1ORF1p in this particular cancer[27]. Here we performed IHC analysis on thirty-nine post-operational oral cancer samples and investigated the expression of L1 ORF1 and ORF2 using our in-house L1ORF1p and L1ORF2p antibodies.The detailed information regarding patients and collected samples are shown in table 1.First, the neoplastic nature of all cancer samples used in this study was confirmed by hematoxylin and eosin staining (Supplementary Figure 3). Our IHC analysis with anti-L1hORF2p showed almost 50% (20 samples) samples were ORF2 positive, and no significant expression was observed in 20 samples (Figure 4, Figure 5 and Supplementary Figure 4). Careful analysis of the IHC data showed that the expression of ORF2p was very high in 12 samples (31%) whereas seven samples (18%) showed a moderate level of expression (Figure 4).TheIHC experiments performed with non-immune mouse serum with anti-His primary antibody didn’t show any signal, and were treated as a negative control (Figure 4). The IHC with GAPDH antibody showed significant expression, and was treated as a positive control (Figure 4)

**Figure 4:**
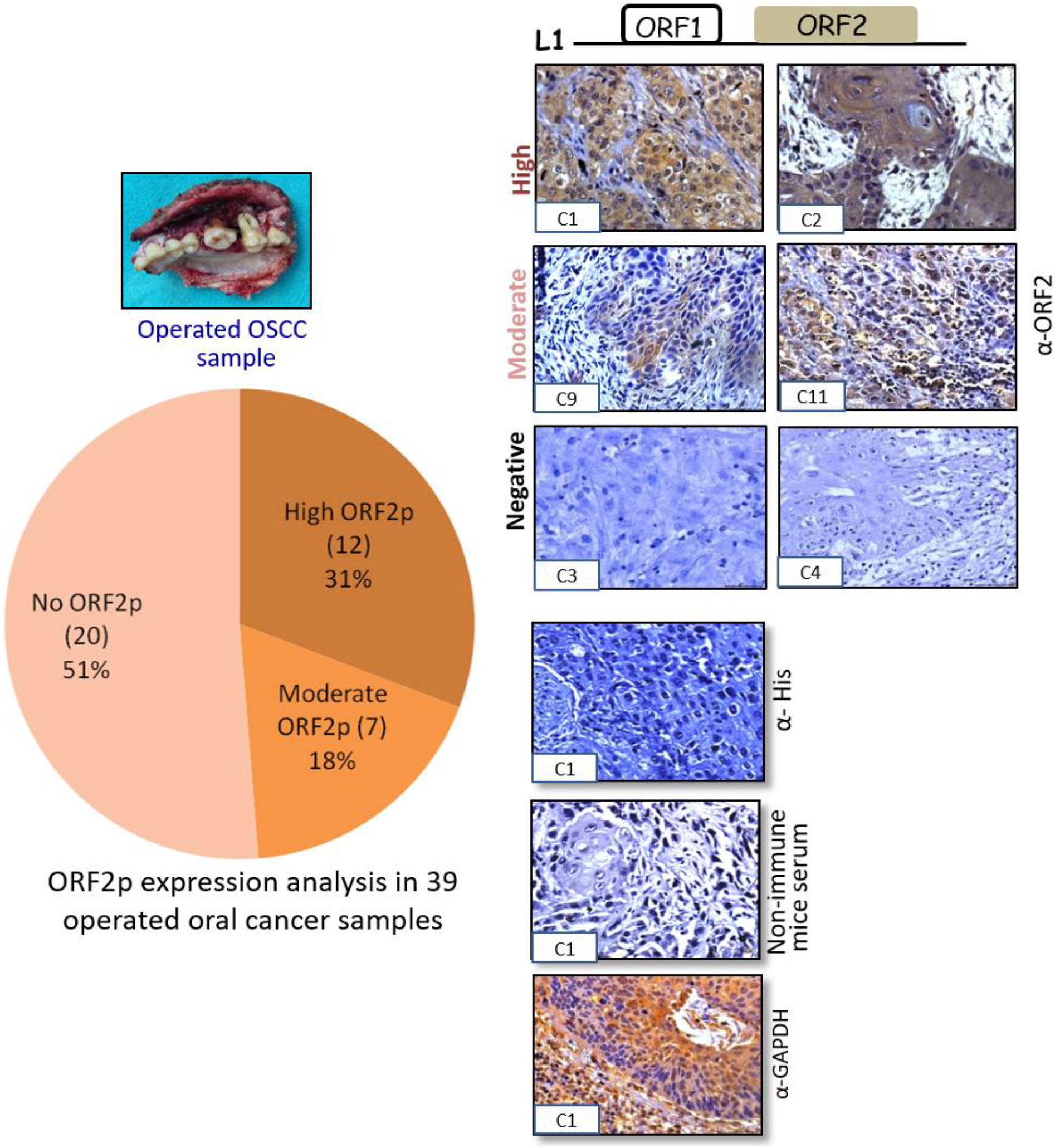
Immuno-peroxidase detection of L1ORF2p expression in post-operative OSCC samples. Immunohistochemistry with anti-L1ORF2p was performed in total 39 post-operativeOSCC. Samples exhibit high (C1 and C2), moderate (C9, C11) and no expression (C3 and C4) (two representatives from each group are shown); IHC staining of the rest of the samples are shown in the supplementary figure.Immuno-staining with anti-His and non-immune mice sera didn’t show any signal. Staining with anti-GAPDH served as a positive control. Images were taken at 40X magnification. The Pie diagram showed almost 50% post-operative OSCC samples expressed a significant amount of L1ORF2p.

**Figure 5:**
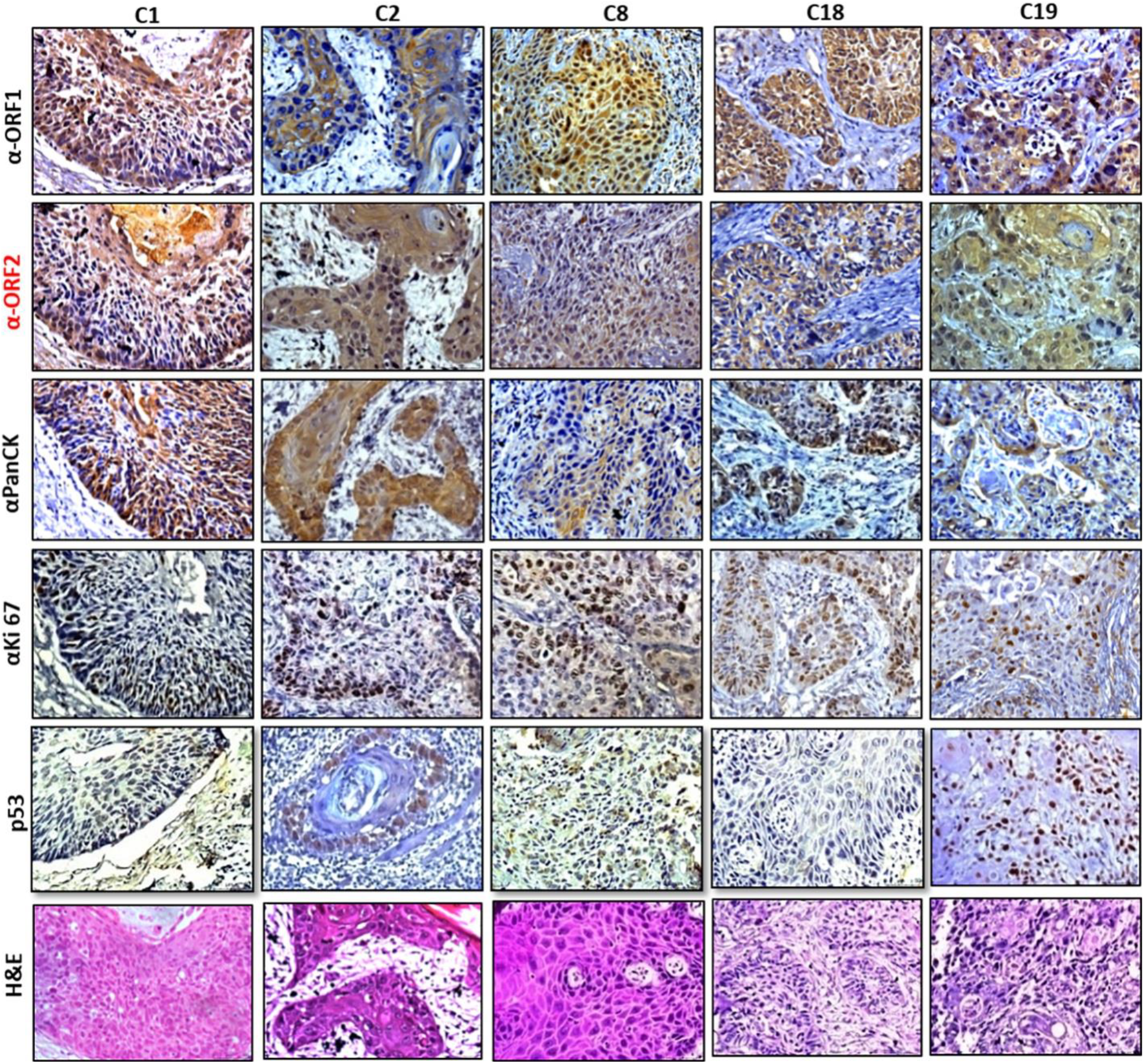
Immuno-peroxidase detection of L1ORF1p, L1ORF2p, Pan-CK, Ki-67 and p53 in operated OSCC samples. IHC staining of five samples (C1, C2, C8, C18, C19) using all five antibodies are shown. The samples were also stained with hematoxylin and eosin.

We next investigated all of the samplesfor expression of hL1ORF1p. The IHC experiment showed that all the twentysamples which were positive for the hL1ORF2p also showed a significant amount of hL1ORF1pexpression(Figure 5A and Supplementary Figure 5).A previous study revealed an increased L1 retrotransposition rate in tumor with p53 mutation [25, 40,41]. Here, we have analysed the aberrant p53 expression by IHC in all the thirty-nine post-operative cancer samples. Our results show significant high expression of p53 in fifteen samples out of thirty sixanalysed(41.6%) (Figure 5A and Supplementary Figure 6). In parallel, the IHC analysis was also performed with anti-PanCK (a marker for epithelial cell) and anti-Ki67 (markers for proliferative cells) for all 39 cancer samples (Figure 5A). The expression of both proteins in all the samples indicated that the post-operational tissues were enriched forcancer cells of epithelial origin.

**Figure 6:**
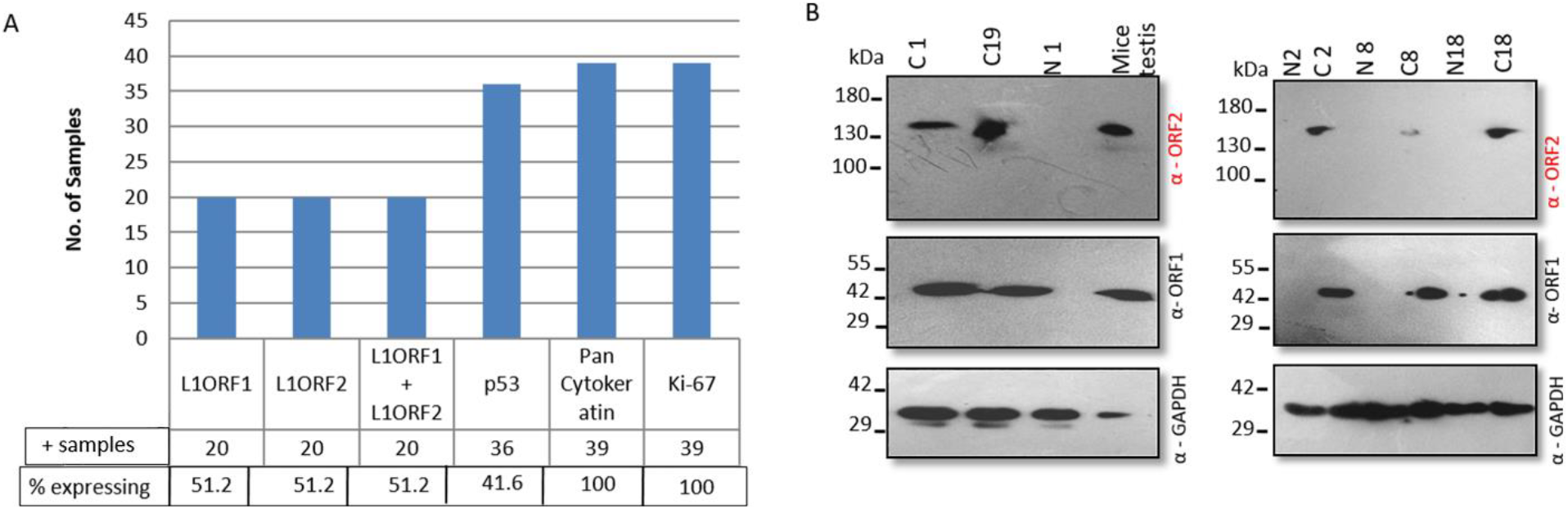
Expression analysis of L1ORF1p and L1ORF2p in OSCC. (A) Bar diagram showing the percent expressing L1ORF1p, L1ORF2p, L1ORF1p + L1ORF2p, p53, Pan-Ck and Ki67. (B) Immunoblot analysis of ORF1p and ORF2p expression in paired normal cancer samples (C1, C2, C8, C18, C19). Anti L1ORF2p detects a distinct band at around 150 kDa corresponds to the MW of ORF2p in all the cancer tissues but not in paired normal. The same blot was re-probed with anti-L1ORF1p and anti-GAPDH.

To complement the IHC results, we performed an immunoblotting experiment to determine the expression of L1ORF1p and L1ORF2p. Three paired samples (normal-cancer pairs), and two samples with only cancer tissues which showed L1-ORFs expression by IHC were tested (Table 1). All five cancer tissues showed a distinct ORF2p band corresponding to150 kDa, the MW of human L1ORF2p. Among the five samples, the sample C19 showed the highest expression, while sample C1 showed the lowest expression of L1ORF2p. Lysate obtained from adjacent normal tissues didn’t show any expression of L1ORF2p. The testis lysate from mice that showed L1ORF2p at around 150 kDawas used as a positive control (Figure 5C).Immunoblot analysis with anti-hL1ORF1p showed very high expression of L1ORF1p in allfive cancer tissues, and no expression was detected in adjacent normal tissues. The GAPDH immunoblotting was used to check the quality of the lysate.

## 4. Discussion

Several studies have reported increased L1 activity in the germ cells andin cancers [22–24,29,39]. In all these studies, our understanding of L1ORF2p expression is limited. Although L1ORF1p is readily detectable [25,27,32], L1ORF2p is very challenging to detect in vitro and in vivo. As the recombinant ORF2p doesn’t express well, short synthetic peptides from the entire length of L1ORF2p protein sequence have been mainly used for the generation of L1ORF2p antibody[28,29, 33–35]. In the past, a number of polyclonal and monoclonal antibodies against human and mouse L1-ORF2p have been reported[28,29,32–35]. However, questions have been raised about the specificity of the reported ORF2p antibodies. Here we found an eighty amino acids stretch at the RT domain [fragment name: hRT_EH_, ORF2p amino acid position 479-558, nucleotide position in L1 RP (Accession No.AF148856) [37] flanked by natural restriction enzymes *EcoRI* and *HindIII* that when cloned and expressed in a bacterial expression vector showed significantly high expression. *In silico* folding of the L1ORF2 protein revealed this particular 80 amino acid stretch protruding outside, making it a suitable epitope for antibody generation. Although the recombinant hL1ORF2RT_EH_fragment peptide formed inclusion bodies, we were successful in purifying and folding it to its native conformation. The purified hL1RTORF2_EH_ fragment induced an adequate antibody response when injected into mice. The whole serum and purified IgG fraction from immunized mice showed a discrete single band at 150 kDa, which corresponds to the MW of hL1ORF2p, suggesting that the antibody is highly specific for detection of L1ORF2 protein.

The eighty amino acids stretch used to make ORF2p antibody showed strong sequence conservation at that particular stretchwhen aligned with mice and rat L1ORF2 sequences. Along with this, we found strong immunoreactivity of this antibody with the mice and rat endogenous L1ORF2p expressed in the germ tissues. Earlier Branciforteet al.[39] reported two mouse-specific ORF2p antibody; both generated using carboxy-terminal two-third of mouse L1-ORF2 protein as antigen. Although the generated antibody showed strong immunoreactivity against the recombinant antigen produced in *E.coli*, no immunoreactivity was detected against endogenous L1ORF2p expressed in mouse testis and ovary. This suggests that the epitope used to generate antibodies against mouse L1ORF2p might be buried inside in the native full-length L1ORF2p, and hence no immunoreactivity was detected [39].

Recently, we have reported a human L1ORF1p antibody which was generated using the RRM domain of human L1ORF1p [26]. Careful observation revealed that amino acid sequences in the RRM domain of the human L1ORF1p are very conserved when compared with mouse and rat L1ORF1 RRM domain amino acids[42,43]. Here we showed that human L1 ORF1p antibody generated against the RRM domain of human L1ORF1p elicitatedastrong immune response against mouse and rat L1ORF1p expressed in the germ tissues. Thus, both the in-house antibodies (anti-hL1ORF1p and anti-hL1ORF2p) detect L1 proteins in all three species (human, mouse and rat) suggesting that these could be useful reagents to study the biology of L1 retrotransposons.

Several studies have demonstrated elevated L1 retrotransposon activity in cancers [23–27,44–47]. L1 promoters, which are heavily methylated in normal tissue to restrict L1 retrotransposition, areoften hypomethylated in tumours leading to L1 retrotransposon activity [48–53]. Whole-genome and targeted sequencing approacheshave showna significant L1 retrotransposon copy number increase in cancer tissues compared to the normal counterpart[23,24,41,45]. Further characterization revealed that the increased copy number of L1 retrotransposons in cancer is the result of active retrotranspositionbythe TPRT mechanism where both the L1 encoded proteins (L1ORF1p and L1ORF2p) are strictly required [24,47]. In a pioneeringstudy, Rodic et al. demonstrated L1ORF1p expression is common, and nearly half of the cancers expressed L1ORF1p as evident by IHC analysis [23]. Attempts to show the L1ORF2 protein expression in cancer tissues are minimal due to the non-availability of an effective L1ORF2p antibody. Here, we have successfully generated a specific L1ORF2p antibody. By employing IHC and immunoblotting, we analysed the expression of L1ORF2p in 39 post-operative oral cancer samples, of which nearly fifty percent samples showed significant L1ORF2p expression.In parallel, we also found L1ORF1p expression in all the samples that showed L1ORF2p expression.Previous studies reported that an increased presence of L1ORF2p in the nucleus is associated with advanced stages of cancer [32,34]. The studies also reported that in some cancers, the expression of L1ORF2p is very high at the early transformation stage[32]. Our data showed among the L1ORF2p positive samples (20 out of 39), some are expressingavery high amount of L1ORF2p. When we compared the amount of ORF2p expression with the grades and TNM staging, no correlation was observed.

The wild type p53 has very short half-life generally localizes in the cell nucleusin very small quantities that it can not be detected by routine IHC[54,55].Mis-sense mutations often increase the half-life and the quantity of p53 expression, allowing its detection by IHC[56]. However, some tumors are frequently immunopositive for p53 in the absence of mutation [57]. Potential mechanisms for accumulation of non-mutant p53 include stabilization of p53protein bybinding to viral or cellular protein and DNA damage by the chemical and physical genotoxic agent [56].

Thus IHC is used a screening test before DNA sequencing to find out missense mutation or overexpression of wild type p53 gene for few cancers. Previous studies showed that in cancer, the increased L1ORF1p expression often correlates with p53 mutations and aberrant p53 expression [25,40]. Wylie et al. [40]demonstrated that the loss of p53 is strongly associated with elevated retrotransposon activity.In the 39 OSCC samples, p53 positive staining was found in 41.6% of the tumors by IHC.Further study is required to under whether mutations in the p53 gene have any role in elevated L1 protein expression in OSCC cancer.

In summary, we have successfully developed a polyclonal L1ORF2p antibody, which is very specific and detects L1ORF2p in post-operative oral cancer tissue. As the epitope used to generate L1ORF2p antibody is highly conserved, we found the antibody is equally useful to detect the same protein in mouseand rat germ tissues.The novel L1ORF2p antibody reported in this study will serve as a useful tool for oral cancer studies and diagnostic applications. Further study is required to understand why L1 activity is deregulated in OSCC and how it contributes to the progression of this particular cancer.

## 5. Conclusions

The L1 retrotransposons show very high activity in germ tissues and many cancers. It expresses two proteins (L1ORF1p and L1ORF2p), both are required for the retrotransposition of L1mRNA. Previous studies demonstrated that almost half of the cancers show the expression of L1ORF1p. However, due to the unavailability of an effective L1ORF2p antibody, the detection of L1ORF2p has not been reported. In this study, we have developed a very specific polyclonal L1ORF2p antibody and showed its robust expression in post-operative oral cancer samples. As the selected epitope used to generate L1ORF2p antibody is highly conserved, we found the antibody is equally useful to detect the same protein in mouse and rat germ tissues. The novel L1ORF2p antibody reported in this study will serve as a useful tool for oral cancer studies and diagnostic applications. Further study is required to understand why L1 activity is deregulated in OSCC and how it contributes to the progression of this particular cancer.

## 6. Author contributions

KM conducted all the experiments and helped to write the manuscript. DS helped in Western, and immunohistochemistry. AS, VKD, BJ and SPA provided all the paired cancer-normal OSCC samples. SR, ND and RS helped in generating the L1ORF2p antibody. SN helped in immunohistochemistry. KVVP helped in immunohistochemistry and analysing data. PKM conceived of the study, supervised experiments, analyzed data and wrote the manuscript.

## 7. Acknowledgements

We thank Dr. Sudha Bhattacharya (School of Environmental Sciences, Jawaharlal Nehru University, New Delhi, India) for helping with reagents and chemicals required in this study. KM, and DSare a recipient of a research fellowship from MHRD, India.

