## Supplementary File for "Robust expression of LINE-1 retrotransposon encoded proteins in oral squamous cell carcinoma"

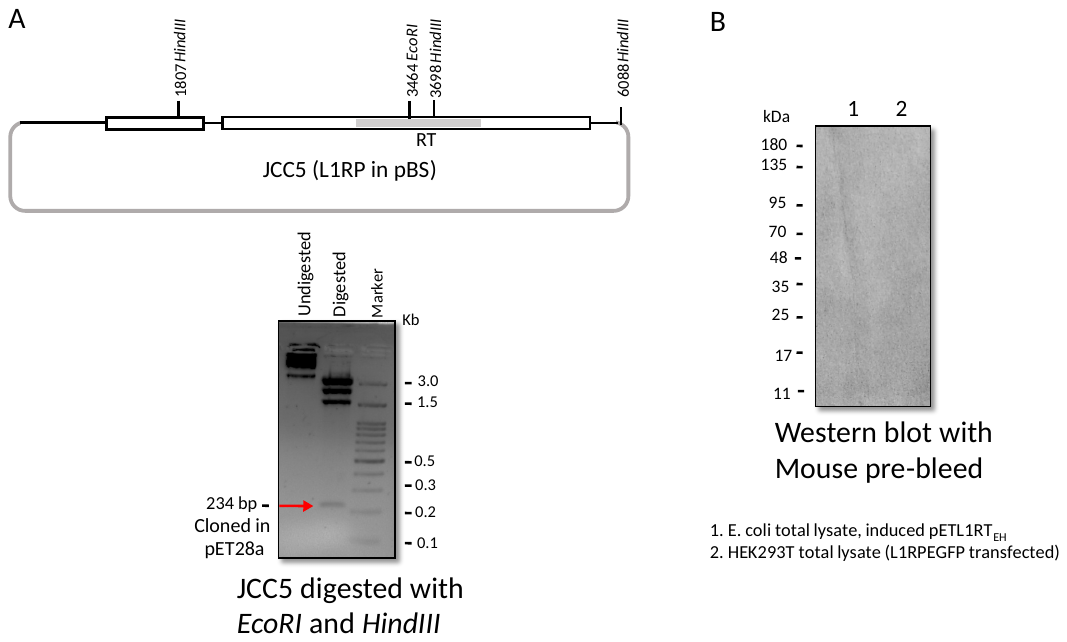

**Supplementary figure 1:** (A) Schematic map of JCC5 (L1RP in pBSKS plasmid) [37] and its restriction digestion with *EcoRI*and*HindIII*restriction enzymes. The digested product was resolved in 1.5% agarose gel. The 234 bp RT fragment (marked by arrow) was subcloned in Pet28a bacterial expression vector. (B) Western blot of L1RT_EH_ (immunogen) and full length exogenous L1ORF2p with non-immune mice sera.

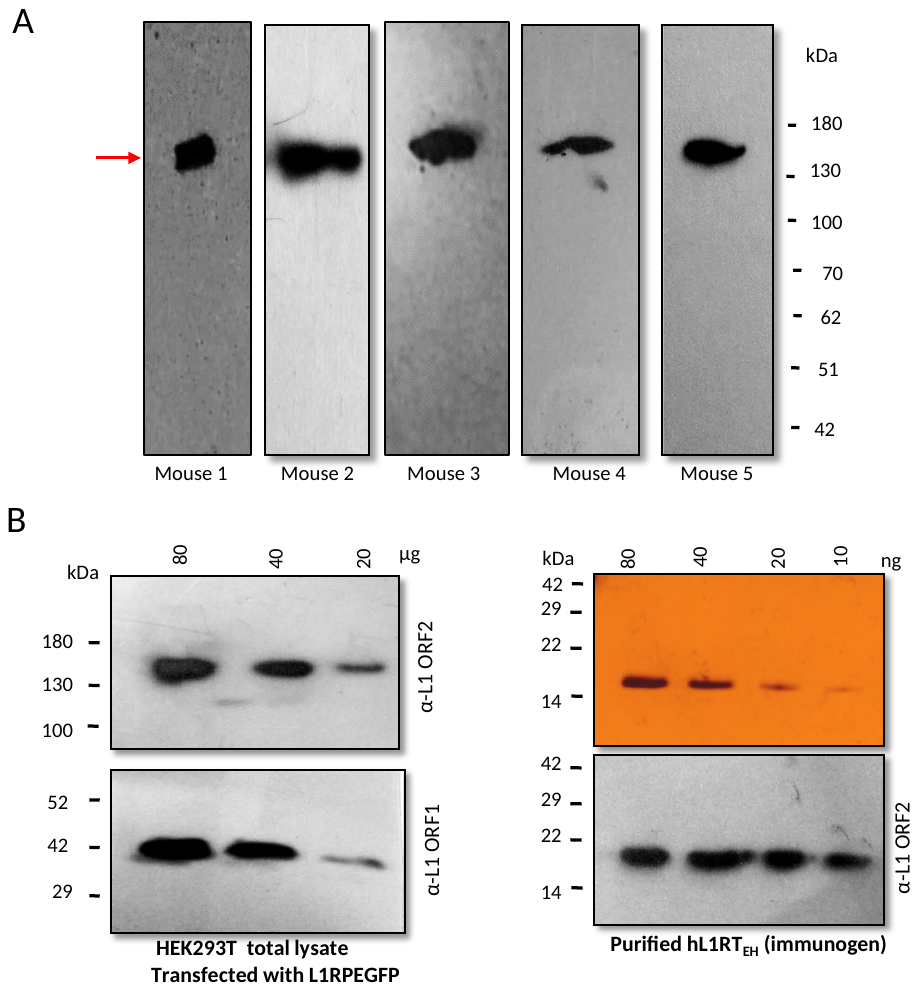

**Supplementary figure 2:** (A) Immune sera from five different mice were checked by Western blot showing a distinct 150 kDa band of exogenous L1ORF2p in HEK293 total lysate transfected with L1RPEGFP [38]. (B) Sensitivity analysis of L1 ORF2p antibody by Western blotting.

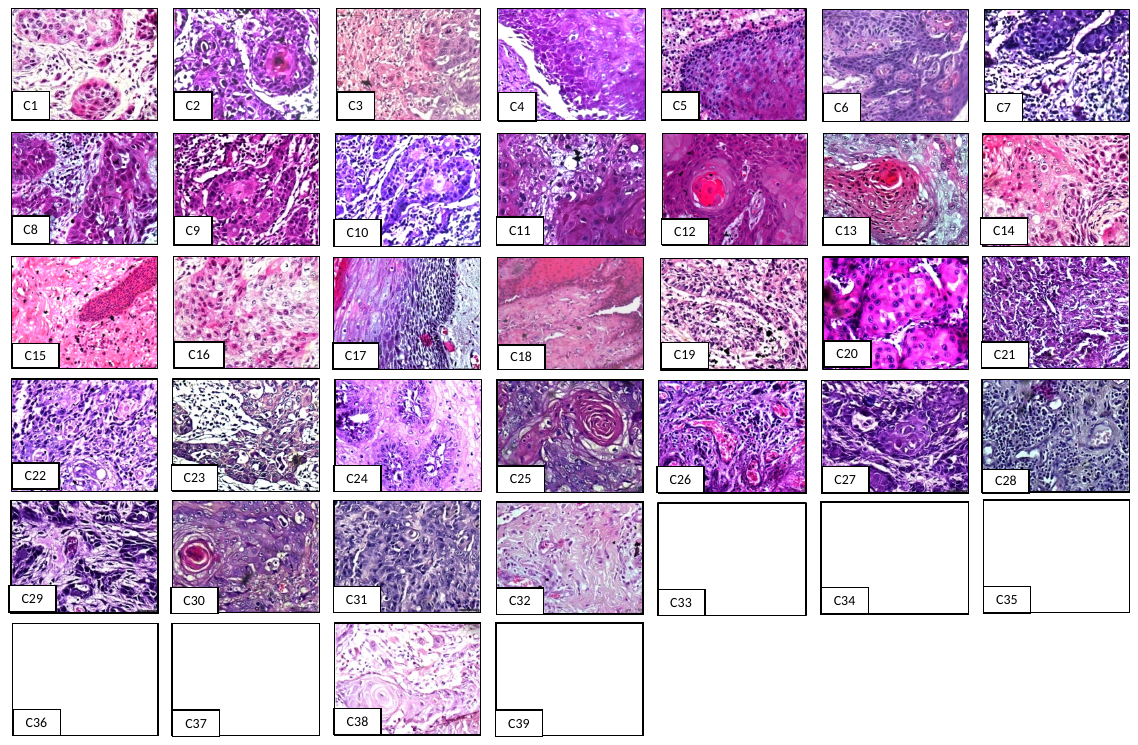

**Supplementary figure 3:** Hematoxylin-eosin stained section of post-operative OSCC samples.

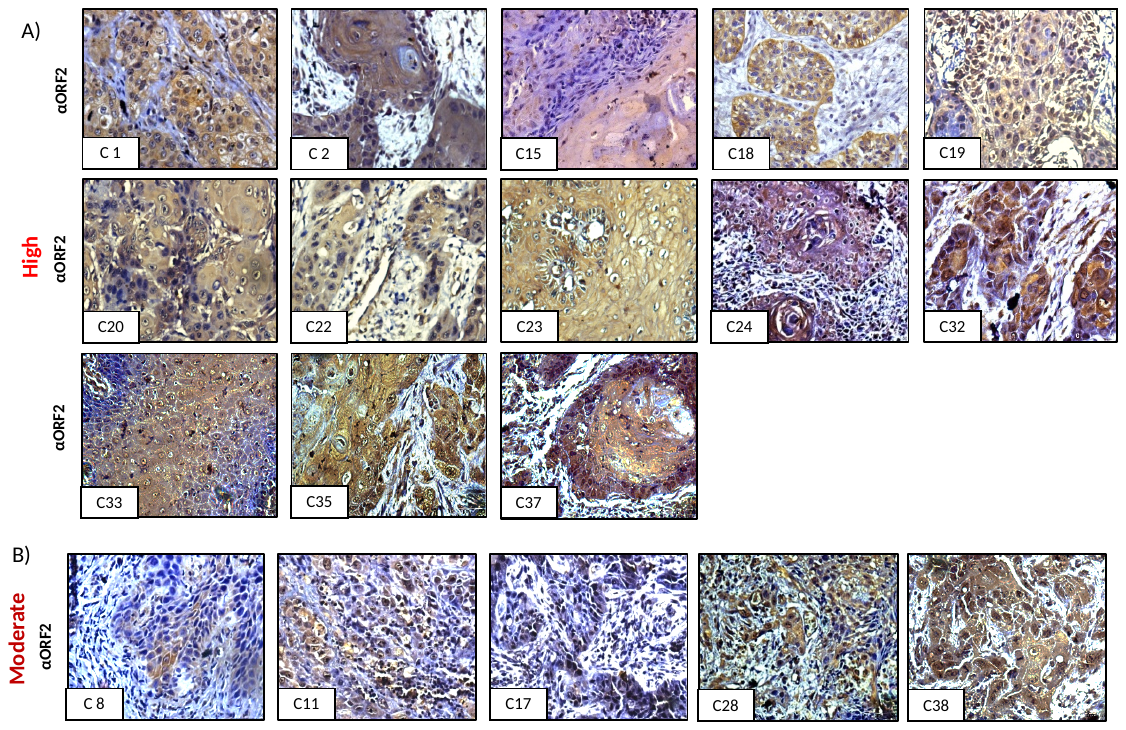

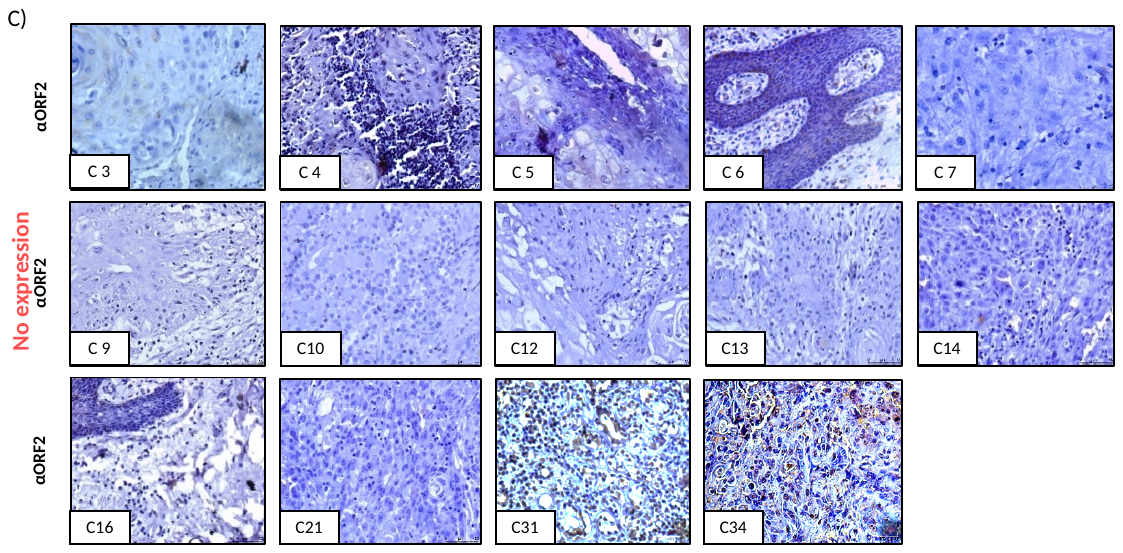

**Supplementary figure 4:** IHC analysis of L1ORF2p expression in post-operative OSCC samples.

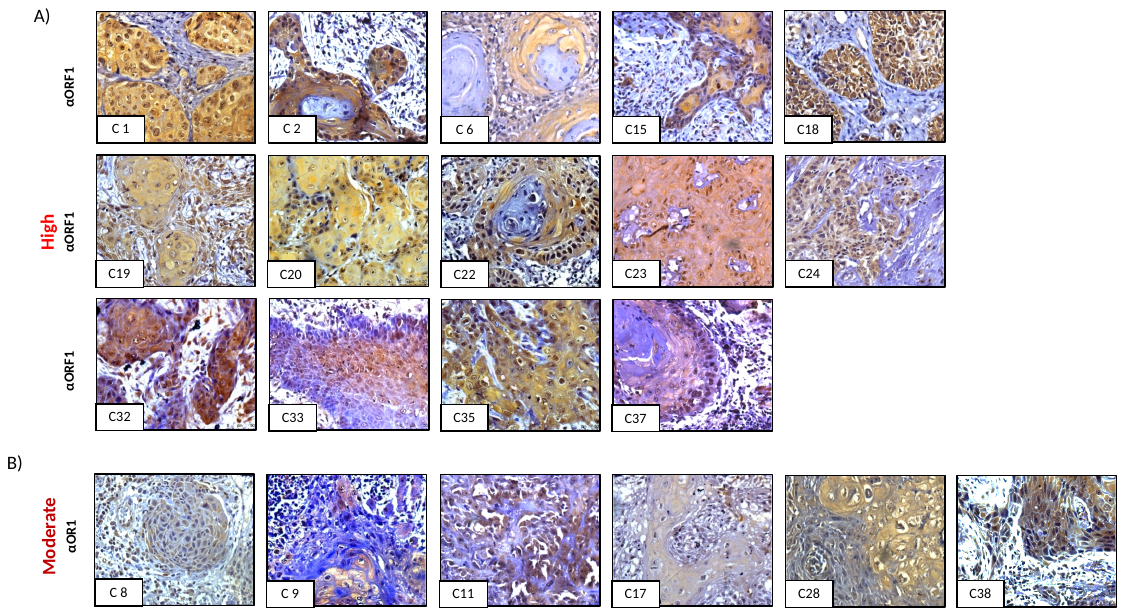

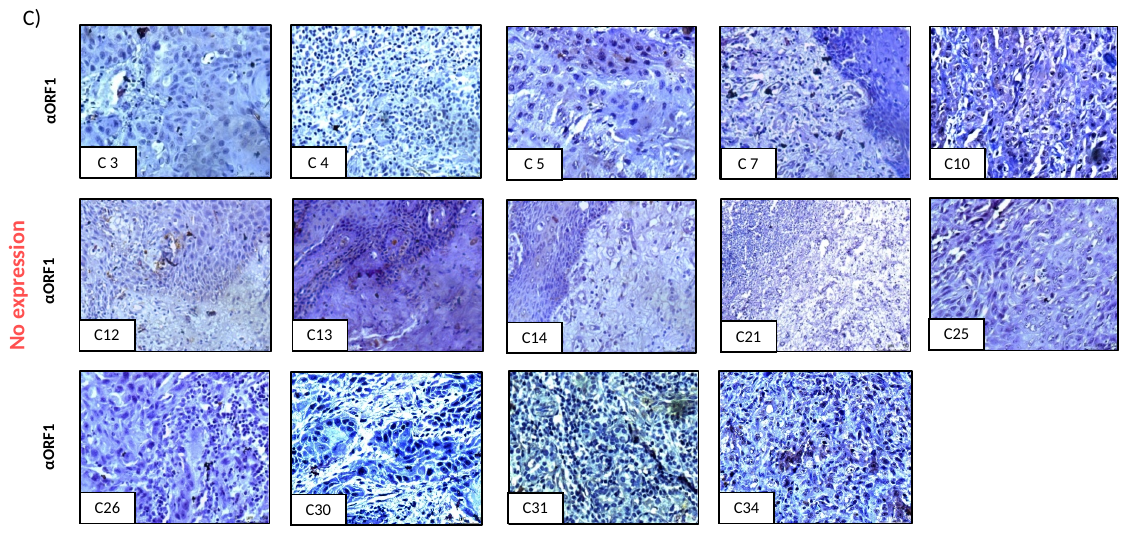

**Supplementary figure 5:** IHC analysis of L1ORF1p expression in post-operative OSCC samples.

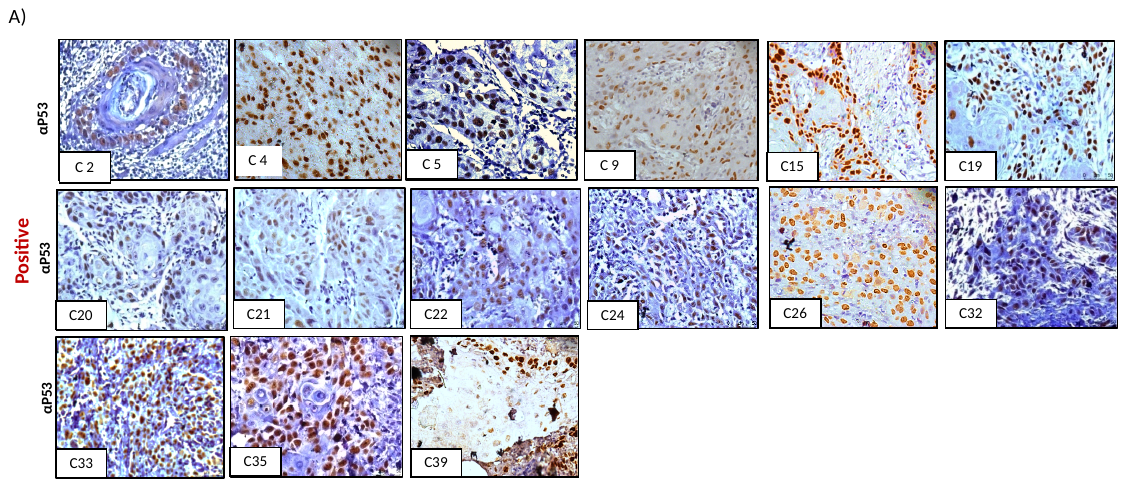

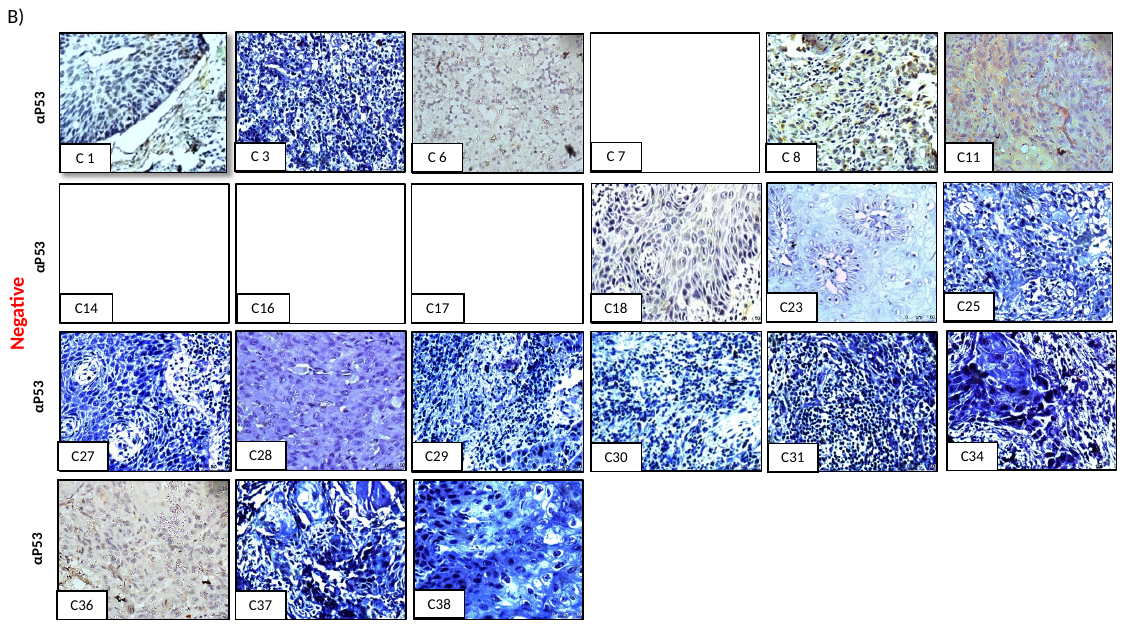

**Supplementary figure 6:** IHC analysis of p53 expression in post-operative OSCC samples.

**Supplementary text:**

**Cloned in pET 28a between ECOR1 and Hind III:**

ATGGGCAGCAGCCATCATCATCATCATCACAGCAGCGGCCTGGTGCCGCGCGGCAGCCATATGGCTAGCATGACTGGTGGACAGCAAATGGGTCGCGGATCCGAATTCTACCAGAGGTACAAGGAGGAACTGGTACCATTCCTTCTGAAACTATTCCAATCAATAGAAAAAGAGGGAATCCTCCCTAACTCATTTTATGAGGCCAGCATCATTCTGATACCAAAGCCGGGCAGAGACACAACCAAAAAAGAGAATTTTAGACCAATATCCTTGATGAACATTGATGCAAAAATCCTCAATAAAATACTGGCAAACCGAATCCAGCAGCACATCAAAAAGCTTGCGGCCGCACTGGAGCACCACCACCACCACCACTGA

Human L1RP (Acc No. AF148856)

Mice L1 (Acc No. M29324.1) (J. Mol. Biol. 196;757-767 ;1987)

Rat L1 (Acc No. DQ100473.1)(Kirilyuk et al. NAR 2008)

>Human L1ORF1 (L1Rp)

MGKKQNRKTGNSKTQSASPPPKERSSSPATEQSWMENDFDELREEGFRRSNYSELREDIQTKGKEVENFEKNLEECITRITNTEKCLKELMELKTKARELREECRSLRSRCDQLEERVSAMEDEMNEMKREGKFREKRIKRNEQSLQEIWDYVKRPNLRLIGVPESDVENGTKLENTLQDIIQENFPNLARQANVQIQEIQRTPQRYSSRRATPRHIIVRFTKVEMKEKMLRAAREKGRVTLKGKPIRLTADLSAETLQARREWGPIFNILKEKNFQPRISYPAKLSFISEGEIKYFIDKQMLRDFVTTRPALKELLKEALNMERNNRYQPLQNHAKM

>Mouse L1 ORF1 (GF21)

MAKGKRKNPTNRSQDHSPSSEPRTPTSPNPGHPNTPEKVDVDLKAYLMMMVEDIKKEFNNSLKEIQENTAKELQVLKEKQENTAKELQVLKEKQENTTKQVEVLIEKQENTSKQVMEMNKTILDLKREVDTIKKTQSEATLEIETLGKKSGTIDASISNRIQEMEERISGAEDSIENIGTTIKENGKCKKILTQNIQEIQDTMRRPNLRIIGVDENEDFQLKGPANIFNKIIEENFPNLKKEMPMNIQEAYRTPNRLDQKRNSSRHIIIRTPNALNKDRILKAVREKGQVTYKGKPIRITPDFSPETMKARRAWTDVIQTLREHKCQPRLLYPAKLSITIDGETKVFHDKTKFTHYLSTNPALQRIITEKKQYKDGNHALEKTRR

>Rat L1 ORF1

MARGKRRNPSNRNQDCMPSSEPNSPAKTNMEYPNTPEKQDLVSKSYLIMMLEDFKKDMNTLRETQEIINKQVEAYREEWQKSLKEFQENTIKQMKELKMEIEAIKKEHMETTLDIENQKKRQGAVDTSFTNRIQEMEERISGAEDSIEIIDSTVKDNVKRKKLLVQNIQEIQDSMRRSNLRIIGIEESEDSQLKGPVNIFNKIIEENFPNLKKEIPIGIQEAYRTPNRLDQKRNTSRHIIVKTPNAQNKERILKAVREKGQVTYKGRPIRITPDFSPETMKARRSWTDVIQTLREHKCQPRLLYPAKLSINIDGETKIFHDKTKFTQYLSTNPALQRIINGKAQHKEASYTLEEARN

CLUSTAL multiple sequence alignment of Human, Mice and Rat L1ORF1 protein sequences.

Yellow shaded part from human L1ORF1 was used to make L1ORF1 specific antibody

```
Human      --MGKKQNRKTGNSKTQSASPPP---KERSSSPATE--QSWMENDFDELREEGFRRSNYS	53
Mouse      MAKGKRKNPTNRSQDHSPSSEPRTPTSPNPGHPNTPEKVDVDLKAYLMMMVEDIKKEFNN	60
Rat        MARGKRRNPSNRNQDCMPSSEPNSPAKTNMEYPNTPEKQDLVSKSYLIMMLEDFKKDMNT	60
              **::* .. ...   :* *    . .   * *    .   : :  :  *.:::.  .

Human      ELREDIQTKGKEVENFE---------------------KNLEECITRITNTEKCLKELME	92
Mouse      SLKEIQENTAKELQVLKEKQENTAKELQVLKEKQENTTKQVEVLIE---KQENTSKQVME	117
Rat        L-RETQEIINKQVEAYREEWQK------SLKE-----------------FQENTI-----	91
             :*  :   *:::  .                                  *:

Human      LKTKARE-------------------------LREECRSLRSRCDQLEERVSAMEDEMNE	127
Mouse      MNKTILDLKREVDTIKKTQSEATLEIETLGKKSGTIDASISNRIQEMEERISGAEDSIEN	177
Rat        --KQMKELKMEIEAIKKEHMETTLDIENQKKRQGAVDTSFTNRIQEMEERISGAEDSIEI	149
             .   :                               *: .* :::***:*. **.::

Human      MKREGKFREKRIKRNEQSLQEIWDYVKRPNLRLIGVPESDVENGTKLENTLQDIIQENFP	187
Mouse      IGTTIKENGKCKKILTQNIQEIQDTMRRPNLRIIGVDENEDFQLKGPANIFNKIIEENFP	237
Rat        IDSTVKDNVKRKKLLVQNIQEIQDSMRRSNLRIIGIEESEDSQLKGPVNIFNKIIEENFP	209
           :    * . *  *   *.:*** * ::* ***:**: *.:  : .   * ::.**:****

Human      NLARQANVQIQEIQRTPQRYSSRRATPRHIIVRFTKVEMKEKMLRAAREKGRVTLKGKPI	247
Mouse      NLKKEMPMNIQEAYRTPNRLDQKRNSSRHIIIRTPNALNKDRILKAVREKGQVTYKGKPI	297
Rat        NLKKEIPIGIQEAYRTPNRLDQKRNTSRHIIVKTPNAQNKERILKAVREKGQVTYKGRPI	269
           ** ::  : ***  ***:* ..:* : ****::  :.  *:::*:*.****:** **:**

Human      RLTADLSAETLQARREWGPIFNILKEKNFQPRISYPAKLSFISEGEIKYFIDKQMLRDFV	307
Mouse      RITPDFSPETMKARRAWTDVIQTLREHKCQPRLLYPAKLSITIDGETKVFHDKTKFTHYL	357
Rat        RITPDFSPETMKARRSWTDVIQTLREHKCQPRLLYPAKLSINIDGETKIFHDKTKFTQYL	329
           *:* *:* **::*** *  ::: *:*:: ***: ******:  :** * * **  : .::

Human      TTRPALKELLKEALNMERNNRYQPLQNHAKM	338
Mouse      STNPALQRIITEKKQYKDGNHALEKTR--R-	385
Rat        STNPALQRIINGKAQHKEASYTLEEAR--N-	357
           :*.***:.::.   : :  .      .  .
```

>Human L1 ORF2

MTGSTSHITILTLNINGLNSAIKRHRLASWIKSQDPSVCCIQETHLTCRDTHRLKIKGWRKIYQANGKQKKAGVAILVSDKTDFKPTKIKRDKEGHYIMVKGSIQQEELTILNIYAPNTGAPRFIKQVLSDLQRDLDSHTLIMGDFNTPLSTLDRSTRQKVNKDTQELNSALHQADLIDIYRTLHPKSTEYTFFSAPHHTYSKIDHIVGSKALLSKCKRTEIITNYLSDHSAIKLELRIKNLTQSRSTTWKLNNLLLNDYWVHNEMKAEIKMFFETNENKDTTYQNLWDAFKAVCRGKFIALNAYKRKQERSKIDTLTSQLKELEKQEQTHSKASRRQEITKIRAELKEIETQKTLQKINESRSWFFERINKIDRPLARLIKKKREKNQIDTIKNDKGDITTDPTEIQTTIREYYKHLYANKLENLEEMDTFLDTYTLPRLNQEEVESLNRPITGSEIVAIINSLPTKKSPGPDGFTAEFYQRYKEELVPFLLKLFQSIEKEGILPNSFYEASIILIPKPGRDTTKKENFRPISLMNIDAKILNKILANRIQQHIKKLIHHDQVGFIPGMQGWFNIRKSINVIQHINRAKDKNHMIISIDAEKAFDKIQQPFMLKTLNKLGIDGTYFKIIRAIYDKPTANIILNGQKLEAFPLKTGTRQGCPLSPLLFNIVLEVLARAIRQEKEIKGIQLGKEEVKLSLFADDMIVYLENPIVSAQNLLKLISNFSKVSGYKINVQKSQAFLYTNNRQTESQIMGELPFTIASKRIKYLGIQLTRDVKDLFKENYKPLLKEIKEETNKWKNIPCSWVGRINIVKMAILPKVIYRFNAIPIKLPMTFFTELEKTTLKFIWNQKRARIAKSILSQKNKAGGITLPDFKLYYKATVTKTAWYWYQNRDIDQWNRTEPSEIMPHIYNYLIFDKPEKNKQWGKDSLFNKWCWENWLAICRKLKLDPFLTPYTKINSRWIKDLNVKPKTIKTLEENLGITIQDIGVGKDFMSKTPKAMATKDKIDKWDLIKLKSFCTAKETTIRVNRQPTTWEKIFATYSSDKGLISRIYNELKQIYKKKTNNPIKKWAKDMNRHFSKEDIYAAKKHMKKCSSSLAIREMQIKTTMRYHLTPVRMAIIKKSGNNRCWRGCGEIGTLLHCWWDCKLVQPLWKSVWRFLRDLELEIPFDPAIPLLGIYPNEYKSCCYKDTCTRMFIAALFTIAKTWNQPKCPTMIDWIKKMWHIYTMEYYAAIKNDEFISFVGTWMKLETIILSKLSQEQKTKHRIFSLIGGN

>Mice L1ORF2

MPTLTTKIKGSNNYFSLISLNINGLNSPIKRHRLTDWLHKQDPTFCCLQETHLREKDRHYLRVKGWKTTFQANGLKKQAGVAILISDKIDFQPKVIKKDKEGHFILIKGKILQEELSILNIYAPNARAATFIRDTLVKLKAYIAPHTIIVGDFNTPLSSKDRSWKQKLNRDTVKLTEVMKQMDLTDIYRAFYPKTKGYTFFSAPHGTFSKIDHIIGHKTGLNRYKNIEIVPCILSDHHGLRLIFNDNINNGKPTFTWKLNNTLFNDTLVKEGIKKEIKDFLEFNENEATTYPNLWDTMKAFLRGKLIALSASKKKRETAHTSSLTTHLKALEKKEAHSPKRSRRQEIIKLRGEINQVETRRTIQRINQTRSWFFEKINKIDKPLARLTKGHRDKILINKIRNEKGDITTDPEEIQNTIRSFYTRLYSTKLENLDEMDKFLDRYQVPKLNQDQVDHLNSPISPKEIEAVINSLPTKKSPGPDGFSAEFYQTFKEDLIPILHKLFHKIEVEGTLPNSFYEATITLIPKPQKDPTKIENFRPISLMNIDAKILNKILANRIQEHIKAIIHPDQVGFIPGMQGWFNIRKSINVIHYINKLKDKNHMIISLDAEKAFDKIQHPFMIKVLERSGIQGPYLNMIKAIYSKPVANIKVNGEKLEAIPLKSGTRQGCPLSPYLFNIVLEVLARAIRQQKEIKGIQIGKEEVKISLFADDMIVYISDPKNSTRELINLINSFGEVAGYKINSNKSMAFLYTKNKQAEKEIRETTPFSIVTNNIKYLGVTLTKEVKDLYDKNFKSLKKEIKEDLRRWKDLPCSWIGRINIVKMAILPKAIYRFNAIPIKIPTQFFNELEGAICKFVWNNKKPRIAKSLLKDKRTSGGITMPDLKLYYRAIVIKTAWYWYRDRQVDQWNRIEDPEMNPHTYGHLIFDKGAKTIQWKKDSIFNNWCWHNWLLSCRRMRIDPYLSPCTKVKSKWIKELHIKPETLKLIEEKVGKSLEDMGTGEKFLNRTAMACAVRSRIDKWDLMKLQSFCKAKDTVNKTKRPPTDWERIFTYPKSDRGLISNIYKELKKVDFRKSNNPIKKWGSELNKEFSPEEYRMAEKHLKKCSTSLIIREMQIKTTLRFHLTPVRMAKIKNSGDSRCWRGCGERGTLLHCWWECRLVQPLWKSVWRFLRKLDIVLPEDPAIPLLGIYPEDAPTGKKDTCSTMFIAALFIIARSWKEPRCPSTEEWIQKMWYIYTMEYYSAIKKNEFMKFLAKWMDLEGIILSEVTHSQRNSHNMYSLISGY

>Rat L1ORF2

MNIKGNNNHYSLISLNINGLNSPIKRHRLTNWIRNEDPAFCCLQETHLRDKDRHYLRVKGWKTTFQANGQKKQAGVAILISNKINFQLKVIKKDKEGHFIFIKGKIHQDELSILNIYAPNTRAPTYVKETLLKLKTHIAPHTIIVGDFNTPLSSMDRSWKQKLNSDVDRLREVMSQMDLTDIYRTFYPKAKGYTFFSAPHGTFSKIDHIIGQKTGLNRYRKIEIIPCVLSDHHGLKLVFNNNKGRMPTYTWKLNNALLNDNLVKEEIKKEIKNFLEFNENENTTYSNLWDTMKAVLRGKLIALSACRKKQERAYVSSMTAHLKALEQKEANTPRRSRRQEIIKLRAEINQVETKRTIERINRTKSWFFEKINKIDKPLARLTRGHRECVQINKIRNEKGDITTDSEEIQKIIRSYYKNLYSTKFENLQEMDYFLDRYQVSKLNQEQINQLNNPITPKEIEAVIKGLPTKKSPGPDGFSAEFYQTFIEDLIPILSKLFHKIETDGALPNSFYESTITLIPKPHKDTTKKGNFRPISLMNIDAKILNKILANRIQEHIKTIIHHDQVGFIPGMQGWFNIRKTINVIHYINKLKEQNHMIISLDAEKAFDKIQHPFMIKVLERIGIQGPYLNIVKAIYSKPVANIKLNGEKLEAIPLKSGTRQGCPLSPYLFNIVLEVLARAIRQQKEIKGMQIGKEEVKISLFADDMIVYLSDPKSSTRELLKLINNFSKVAGYKINSNKSVAFLYTKEKQAEKEIRETTPFIIDPNNIKYLGVALTKQVKDLYNKNFKTLKKEIEEDLRRWKDLPCSWIGRINIVKMAILPKAIYRFNAIPIKIPIQFFKELDRTICKFIWNNKKPRIAKAILNNKRTSGGITIPELKQYYRAIVIKTAWYWYRDRQIDQWNRIEDPEMNPHTYGHLIFDKGAKTIQWKKDSIFSKWCWFNWRATCRRMQIDPCLSPCTKLKSKWIKDLHIKPDTLKLIEEKLGKHLEHMGTGKNFLNKTPMAYALRSRIDKWDLIKLQSFCKAKDTVVRTKRQPTDWEKIFTNPTTDRGLISKIYKELKKLDRRETNNPIKKWGSELNKEFTAEECRMAEKHLKKCSTSLVIREMQIKTTLRFHLTPVRLAKIKNSGDSRCWRGCGERGTLLHCWWDCRLVKPFWKSVWRFLRKLDIELPEDPAIPLLGIYPKDASTYKRDTCSTMFIAALFIIARKWKEPRCPSTEEWIQKMWYIYTMEYYSAIKNNKFMKFVGKWLELENIILSELTQSQKDIHGMHSLISGY

CLUSTAL multiple sequence alignment of Human, Mice and Rat L1ORF2 protein sequences

Yellow shaded part from human L1ORF2 was used to make L1ORf2 specific antibody

```
Human_ORF2      -------MTGSTSHITILTLNINGLNSAIKRHRLASWIKSQDPSVCCIQETHLTCRDTHR	53
ORF2_GF21       MPTLTTKITGSNNYLSLISLNINGLNSPIKRHRLTDWLHKQDPTFCCLQETHLREKDRHY	60
RAT_ORF2        -----MNIKGNNNHYSLISLNINGLNSPIKRHRLTNWIRNEDPAFCCLQETHLRDKDRHY	55
                       :.*...: ::::******** ******:.*::.:**:.**:*****  :* *

Human_ORF2      LKIKGWRKIYQANGKQKKAGVAILVSDKTDFKPTKIKRDKEGHYIMVKGSIQQEELTILN	113
ORF2_GF21       LRVKGWKTIFQANGLKKQAGVAILISDKIDFQPKVIKKDKEGHFILIKGKILQEELSILN	120
RAT_ORF2        LRVKGWKTTFQANGQKKQAGVAILISNKINFQLKVIKKDKEGHFIFIKGKIHQDELSILN	115
                *::***:. :**** :*:******:*:* :*: . **:*****:*::**.* *:**:***

Human_ORF2      IYAPNTGAPRFIKQVLSDLQRDLDSHTLIMGDFNTPLSTLDRSTRQKVNKDTQELNSALH	173
ORF2_GF21       IYAPNARAATFIKDTLVKLKAHIAPHTIIVGDLNTPLSSMDRSWKQKLNRDTVKLTEVMK	180
RAT_ORF2        IYAPNTRAPTYVKETLLKLKTHIAPHTIIVGDFNTPLSSMDRSWKQKLNSDVDRLREVMS	175
                *****: *  ::*:.* .*: .:  **:*:**:*****::*** :**:* *. .* ..:

Human_ORF2      QADLIDIYRTLHPKSTEYTFFSAPHHTYSKIDHIVGSKALLSKCKRTEIITNYLSDHSAI	233
ORF2_GF21       QMDLTDIYRIFNPKTKGYTFFSAPHGTFSKIDHIIGHKTGLNRYKNIEIVPCILSDHHGL	240
RAT_ORF2        QMDLTDIYRTFYPKAKGYTFFSAPHGTFSKIDHIIGQKTGLNRYRKIEIIPCVLSDHHGL	235
                * ** **** : **:. ******** *:******:* *: *.: :. **:   **** .:

Human_ORF2      KLELRIKNLTQSRSTTWKLNNLLLNDYWVHNEMKAEIKMFFETNENKDTTYQNLWDAFKA	293
ORF2_GF21       RLIFNNNIKNGKPTFTWKLNNTLLNDTLVKEGIKKEIKDFLEFNENEATTYPNLWDTMKA	300
RAT_ORF2        KLVFNNN-KGRMPTYTWKLNNALLNDNLVKEEIKKEIKNFLEFNENENTTYSNLWDTMKA	294
                :* :. :      : ****** ****  *:: :* *** *:* ***: *** ****::**

Human_ORF2      VCRGKFIALNAYKRKQERSKIDTLTSQLKELEKQEQTHSKASRRQEITKIRAELKEIETQ	353
ORF2_GF21       FLRGKLIALSTSKKKRERAHTSSLTTHLKALEKKEANSPKRSRRQEIIKLRGEINQVETR	360
RAT_ORF2        VLRGKLIALSACRKKQERAYVSSMTAHLKALEQKEANTPRRSRRQEIIKLRAEINQVETK	354
                . ***:***.: ::*:**:  .::*::** **::* .  : ****** *:*.*::::**:

Human_ORF2      KTLQKINESRSWFFERINKIDRPLARLIKKKREKNQIDTIKNDKGDITTDPTEIQTTIRE	413
ORF2_GF21       RTIQRINQTRSWFFEKINKIDKHLARLTRGQRDKILINKIRNEKGDITTDPEEIQNTIRS	420
RAT_ORF2        RTIERINRTKSWFFEKINKIDKPLARLTRGHRECVQINKIRNEKGDITTDSEEIQKIIRS	414
                :*:::**.::*****:*****: **** : :*:   *:.*:*:*******  ***. **.

Human_ORF2      YYKHLYANKLENLEEMDTFLDTYTLPRLNQEEVESLNRPITGSEIVAIINSLPTKKSPGP	473
ORF2_GF21       FYKSLYSTKLENLDEMDKFLDKYQVPKLNQDQVDLLNSPISPKEIEAVINSLPAKKSPGP	480
RAT_ORF2        YYKNLYSTKFENLQEMDYFLDRYQVSKLNQEQINQLNNPITPKEIEAVIKGLPTKKSPGP	474
                :** **:.*:***:*** *** * : :***:::: ** **: .** *:*:.**:******

Human_ORF2      DGFTAEFYQRYKEELVPFLLKLFQSIEKEGILPNSFYEASIILIPKPGRDTTKKENFRPI	533
ORF2_GF21       DGFSAEFYQTFKEDLIPVLHKLFHRIEVEGTLPNSFYEATITLIPKPQKDPTKIENFRPI	540
RAT_ORF2        DGFSAEFYQTFIEDLIPILSKLFHKIETDGALPNSFYESTITLIPKPHKDTTKKGNFRPI	534
                ***:***** : *:*:*.* ***: ** :* *******::* ***** :* **  *****

Human_ORF2      SLMNIDAKILNKILANRIQQHIKKLIHHDQVGFIPGMQGWFNIRKSINVIQHINRAKDKN	593
ORF2_GF21       SLMNIDAKILNKILANRIQEHIKEIIHPDQVGFIPGMQGWFNIRKSINVIHYINKLKDKN	600
RAT_ORF2        SLMNIDAKILNKILANRIQEHIKTIIHHDQVGFIPGMQGWFNIRKTINVIHYINKLKEQN	594
                *******************:*** :** *****************:****::**: *::*

Human_ORF2      HMIISIDAEKAFDKIQQPFMLKTLNKLGIDGTYFKIIRAIYDKPTANIILNGQKLEAFPL	653
ORF2_GF21       HMIISLDAEKAFDKIQHPFMIKVLERSGIQGPYLNIIKAIYSKPVANIKVNGEKLEAIPL	660
RAT_ORF2        HMIISLDAEKAFDKIQHPFMIKVLERIGIQGPYLNIVKAIYSKPVANIKLNGEKLEAIPL	654
                *****:**********:***:*.*:: **:* *::*::***.**.*** :**:****:**

Human_ORF2      KTGTRQGCPLSPLLFNIVLEVLARAIRQEKEIKGIQLGKEEVKLSLFADDMIVYLENPIV	713
ORF2_GF21       KSGTRQGCPLSPYLFNIVLEVLARAIRQQKEIKGIQIGKEEVKISLFADDMIVYISDPKN	720
RAT_ORF2        KSGTRQGCPLSPYLFNIVLEVLARAIRQQKEIKGMQIGKEEVKISLFADDMIVYLSDPKS	714
                *:********** ***************:*****:*:******:**********:.:*

Human_ORF2      SAQNLLKLISNFSKVSGYKINVQKSQAFLYTNNRQTESQIMGELPFTIASKRIKYLGIQL	773
ORF2_GF21       STRELLNLINSFGEVAGYKINSNKSMAFLYTKNKQAEKEIRETTPFSIVTNNIKYLGVTL	780
RAT_ORF2        STRELLKLINNFSKVAGYKINSNKSVAFLYTKEKQAEKEIRETTPFIIDPNNIKYLGVAL	774
                *:::**:**..*.:*:***** :** *****:::*:*.:*    ** *  :.*****: *

Human_ORF2      TRDVKDLFKENYKPLLKEIKEETNKWKNIPCSWVGRINIVKMAILPKVIYRFNAIPIKLP	833
ORF2_GF21       TKEVKDLYDKNFKSLKKEIKEDLRRWKDLPCSWIGRINIVKMAILPKAIYRFNAIPIKIP	840
RAT_ORF2        TKQVKDLYNKNFKTLKKEIEEDLRRWKDLPCSWIGRINIVKMAILPKAIYRFNAIPIKIP	834
                *::****:.:*:* * ***:*: .:**::****:*************.**********:*

Human_ORF2      MTFFTELEKTTLKFIWNQKRARIAKSILSQKNKAGGITLPDFKLYYKATVTKTAWYWYQN	893
ORF2_GF21       TQFFNELEGAICKFIWNNKKPRIAKTLLKDKRTSGGITMPDLKLYYRAIVIKTAWYWYRD	900
RAT_ORF2        IQFFKELDRTICKFIWNNKKPRIAKAILNNKRTSGGITIPELKQYYRAIVIKTAWYWYRD	894
                  **.**: :  *****:*: ****::*.:*..:****:*::* **:* * *******::

Human_ORF2      RDIDQWNRTEPSEIMPHIYNYLIFDKPEKNKQWGKDSLFNKWCWENWLAICRKLKLDPFL	953
ORF2_GF21       RQVDQWNRIEDPEMNPHTYGHLIFDKGAKTIQWKKDSIFNNWCWHNWLLSCRRMRIYPYL	960
RAT_ORF2        RQIDQWNRIEDPEMNPHTYGHLIFDKGAKTIQWKKDSIFSKWCWFNWRATCRRMQIDPCL	954
                *::***** *  *: ** *.:*****  *. ** ***:*.:*** **   **:::: * *

Human_ORF2      TPYTKINSRWIKDLNVKPKTIKTLEENLGITIQDIGVGKDFMSKTPKAMATKDKIDKWDL	1013
ORF2_GF21       SPCTKVKSKWIKELHIKPETLKLIEEKVGKSLEDMGTGEKFLNRTALACSVRSRIDKWDL	1020
RAT_ORF2        SPCTKLKSKWIKDLHIKPDTLKLIEEKLGKHLEHMGTGKNFLNKTPMAYALRSRIDKWDL	1014
                :* **::*:***:*::**.*:* :**::*  ::.:*.*:.*:.:*  * : :.:******

Human_ORF2      IKLKSFCTAKETTIRVNRQPTTWEKIFATYSSDKGLISRIYNELKQIYKKKTNNPIKKWA	1073
ORF2_GF21       MKLQSFCKAKDTVNKTKRPPTDWERIFTYPKSDRGLISNIYKELKKVDFRKSNNPIKKWG	1080
RAT_ORF2        IKLQSFCKAKDTVVRTKRQPTDWEKIFTNPTTDRGLISKIYKELKKLDRRETNNPIKKWG	1074
                :**:***.**:*. :.:* ** **:**:  .:*:****.**:***::  :::*******.

Human_ORF2      KDMNRHFSKEDIYAAKKHMKKCSSSLAIREMQIKTTMRYHLTPVRMAIIKKSGNNRCWRG	1133
ORF2_GF21       SELNKEFSPEEYRMAEKHLKKCSTSLIIREMQIKTTLRFHLTPVRMAKIKNSGDSRCWRG	1140
RAT_ORF2        SELNKEFTAEECRMAEKHLKKCSTSLVIREMQIKTTLRFHLTPVRLAKIKNSGDSRCWRG	1134
                .::*:.*: *:   *:**:****:** *********:*:******:* **:**:.*****

Human_ORF2      CGEIGTLLHCWWDCKLVQPLWKSVWRFLRDLELEIPFDPAIPLLGIYPNEYKSCCYKDTC	1193
ORF2_GF21       CGERGTLLHCWWDCRLVQPLWKSVWRFLRKLDIVLPEDPAIPLLGIYPEDAP-TGKKDTC	1199
RAT_ORF2        CGERGTLLHCWWDCRLVKPFWKSVWRFLRKLDIELPEDPAIPLLGIYPKDAS-TYKRDTC	1193
                *** **********:**:*:*********.*:: :* ***********::      :***

Human_ORF2      TRMFIAALFTIAKTWNQPKCPTMIDWIKKMWHIYTMEYYAAIKNDEFISFVGTWMKLETI	1253
ORF2_GF21       STMFIAALFIIARSWKEPRCPSTEEWIQKMWYIYTMEYYSAIKKNEFMKFLAKWMDLEGI	1259
RAT_ORF2        STMFIAALFIIARKWKEPRCPSTEEWIQKMWYIYTMEYYSAIKNNKFMKFVGKWLELENI	1253
                : ******* **:.*::*:**:  :**:***:*******:***:::*:.*:..*:.** *

Human_ORF2      ILSKLSQEQKTKHRIFSLIGGN	1275
ORF2_GF21       ILSEVTHSQRNSHNMYSLISGY	1281
RAT_ORF2        ILSELTQSQKDIHGMHSLISGY	1275
                ***::::.*:  * :.***.*
```

>hL1ORF2RT

EFYQRYKEELVPFLLKLFQSIEKEGILPNSFYEASIILIPKPGRDTTKKENFRPISLMNIDAKILNKILANRIQQHIKKL

>mL1ORF2RT

EFYQTFKEDLIPVLHKLFHRIEVEGTLPNSFYEATITLIPKPQKDPTKIENFRPISLMNIDAKILNKILANRIQEHIKEI

>rL1ORF2RT

EFYQTFIEDLIPILSKLFHKIETDGALPNSFYESTITLIPKPHKDTTKKGNFRPISLMNIDAKILNKILANRIQEHIKTI

```
CLUSTAL(1.2.4) multiple sequence alignment


hL1ORF2RT      EFYQRYKEELVPFLLKLFQSIEKEGILPNSFYEASIILIPKPGRDTTKKENFRPISLMNI	60
mL1ORF2RT      EFYQTFKEDLIPVLHKLFHRIEVEGTLPNSFYEATITLIPKPQKDPTKIENFRPISLMNI	60
rL1ORF2RT      EFYQTFIEDLIPILSKLFHKIETDGALPNSFYESTITLIPKPHKDTTKKGNFRPISLMNI	60
               **** : *:*:*.* ***: ** :* *******::* ***** :* **  **********

hL1ORF2RT      DAKILNKILANRIQQHIKKL	80
mL1ORF2RT      DAKILNKILANRIQEHIKEI	80
rL1ORF2RT      DAKILNKILANRIQEHIKTI	80
               **************:*** :


Percent Identity  Matrix - created by Clustal2.1
     1: hL1ORF2RT   100.00   76.25   73.75
     2: mL1ORF2RT    76.25  100.00   83.75
     3: rL1ORF2RT    73.7583.75  100.00
```
